# Early warning signals do not predict a warming-induced experimental epidemic

**DOI:** 10.1101/2025.04.14.648789

**Authors:** Madeline Jarvis-Cross, Devin Kirk, Leila Krichel, Pepijn Luijckx, Péter K. Molnár, Martin Krkošek

**Author notes:** Corresponding author: Madeline Jarvis-Cross. **Open Research Statement:** Empirical and simulated time series data, and scripts available from GitHub at https://github.com/MadelineJC/EarlyWarningSignals_Repo.

## Abstract

Climate change can impact the rates at which parasites are transmitted between hosts, ultimately determining if and when an epidemic will emerge. As such, our ability to predict climate-mediated epidemic emergence will become increasingly important in our efforts to prepare for and mitigate the effects of disease outbreaks on ecological systems and human health. Theory suggests that statistical signatures termed “early warning signals” (EWS), can function as predictors of disease emergence. Here, we analyze experimental and simulated time series of disease spread within populations of the model disease system *Daphnia magna*–*Ordospora colligata* for EWS of epidemic emergence. In this system, low temperatures prevent disease emergence, while sufficiently high temperatures force the system through a critical transition to an epidemic state. We found that EWS of epidemic emergence were nearly as likely to be detected in populations maintained at a sub-epidemic temperature as they were to be detected in populations subjected to a warming treatment that induced epidemic spread. Our findings suggest that the detection of false positives may limit the reliability of EWS as predictors of climate-mediated epidemic emergence.

## Introduction

Host-parasite dynamics are often mediated by temperature. As such, climate change can impact the rates at which parasites are transmitted between hosts, and ultimately, affect if and when epidemics emerge (Altizer et al., 2013; Daszak et al., 2001; Lafferty, 2009; Patz et al., 2008). For example, climate change can alter species ranges and population densities, thereby altering contact rates, and thus, parasite transmission rates (Baker et al., 2022; Rohr et al., 2011). In vector-borne disease systems like malaria and dengue fever, climate change may increase vector range and competence, though subsequent impacts on incidence and epidemic emergence have been debated (Caminade et al., 2014; Childs et al., 2024; Lafferty & Mordecai, 2016). Climate change can also asymmetrically affect vital rates within hosts and their parasites, widening performance gaps between them, and changing infection patterns (Cohen et al., 2017, 2020). In amphibians and a variety of marine organisms (e.g. shellfish, corals, and finfish), climate change has been linked to decreased host immune function and/or increased pathogen virulence, resulting in widespread disease-mediated declines (Burge et al., 2014; Rohr et al., 2008; Rohr & Raffel, 2010). In contrast, increasing mean temperatures have also been associated with decreased infectivity of cold-water pathogens in salmonids (Holt et al., 1989). In general, a wide array of studies of terrestrial and aquatic systems have shown system-specific impacts of climate change on disease spread and emergence (Altizer et al., 2013; Hall et al., 2006; Harvell et al., 2002). As Earth’s climate changes, our ability to predict climate-mediated epidemic emergence will become increasingly important in our efforts to prepare for and mitigate the effects of disease outbreaks on ecological systems, agriculture, and human health. It has been proposed that such predictions may be made by leveraging the statistical signatures associated with ecological time series data.

As dynamical systems, ecological systems are often characterized by critical thresholds (Scheffer, 2009; Scheffer et al., 2001). When parameters that define a system change, the system may pass through a local bifurcation and experience a critical transition, whereby the stability of an equilibrium changes, or the system quickly shifts among alternative stable states. As a system approaches a local bifurcation, its resilience decreases, and the rate at which it recovers from small perturbations slows down (termed “critical slowing down”) (Scheffer et al., 2009; Van Nes & Scheffer, 2007; Wissel, 1984). Theory suggests that as a system’s resilience decreases, it will produce statistically anomalous behaviors, such as increasing variation and temporal autocorrelation. These statistical signatures have been termed “early warning signals” (EWS), and, when detected, have been employed as a means of predicting upcoming critical transitions (Scheffer, 2009; Scheffer et al., 2009).

Although contentious (Boettiger & Hastings, 2012; Dakos et al., 2015), the detection of EWS has been used to assert the predictability of critical transitions in a broad range of dynamical systems, including climate systems (Bury et al., 2021; Dakos et al., 2008; Lenton, 2011; Lenton et al., 2017), freshwater systems (D’Souza et al., 2015; Gsell et al., 2016; O’Brien et al., 2023; Rohde et al., 2022), and infectious disease systems (Dablander et al., 2022; Drake & Hay, 2017; Harris et al., 2020; Miller et al., 2017; O’Regan et al., 2016; O’Regan & Drake, 2013; Pananos et al., 2017). In epidemiology, when the basic reproductive number (R_0_) exceeds a value of one, infectious disease systems pass through a transcritical bifurcation, and shift from a sub-epidemic state to an epidemic state (Drake et al., 2019; O’Regan & Drake, 2013). As such, EWS have been employed to analyze prevalence time series of past epidemics to determine if epidemic (re-)emergence could have been predicted (Southall et al., 2021). For example, Harris et al. (2020) analyzed long-term malaria incidence data and detected increases in eight of ten EWS in the period preceding epidemic emergence. While studies of different disease systems have yielded varying results (Southall et al., 2021), the absence of “control” populations within which sub-epidemic spread fails to transition into epidemic spread limits our ability to interpret these results. In addition, it has been posited that the study of observed systems known to exhibit critical transitions introduces a kind of selection bias, whereby the likelihood of observing evidence of a phenomenon is confused for the likelihood of the phenomenon, given the evidence (Boettiger & Hastings, 2012).

Building off of Kirk et al. (2020), we analyzed experimental and simulated time series of disease spread within populations of the model disease system *Daphnia magna*–*Ordospora colligata* for EWS of critical transition to an epidemic state. During Kirk et al.’s (2020) experiment, control populations were maintained at a sub-epidemic temperature and test populations were subjected to a warming treatment that forced the system through a critical transition to an epidemic state. We found that EWS of epidemic emergence were nearly as likely to be detected in control populations as they were to be detected in warming populations, implying that the ubiquity of false positive detections may limit the reliability of EWS as predictors of climate-mediated epidemic emergence.

## Methods

First, we will describe Kirk et al.’s (2020) experiment and the resulting population level infection data. Then, we will describe Kirk et al.’s (2020) trait-based mechanistic model of O. colligata transmission dynamics, and how we used this model to simulate time series of disease spread under constant and warming conditions. Finally, we will describe how we analysed the experimental and simulated data for early warning signals of disease emergence, and four ways in which we evaluated the reliability of our empirical detections.

### Warming Experiment

The empirical data were derived from a 120-day experiment involving eight *Daphnia magna* populations and their environmentally transmitted microsporidian parasite, *Ordospora colligata* (Kirk et al., 2020b). *D. magna* is a freshwater planktonic crustacean often used as a model organism to develop and evaluate ecological theories (Ebert, 2005). Since the early 1990s, *D. magna* has been used to study host–parasite dynamics at the within- and between-host levels (Ebert, 2005). In nature, *D. magna* hosts numerous parasites, including *O. colligata*, a microsporidian parasite that infects *D. magna*’s gut epithelial cells (Larsson et al., 1997).

Prior to the beginning of the experiment, uninfected *D. magna* were maintained at 20°C for fifteen days, and subsequently, 10°C for an additional fifteen days to acclimatize the populations to the starting conditions of the experiment. Population abundances stabilized between 150 and 240 large individuals per population. The experiment was initiated by introducing three randomly selected adult *D. magna* from infected stock populations to each of the eight experimental populations. Disease prevalence in infected stock populations was approximately 46.5%. Every third day of the experiment, twelve large females were removed from each experimental population and assessed for infection. Sampled individuals were then replaced by three randomly selected individuals from infected stock populations, and nine individuals from uninfected stock populations to maintain an immigration rate of infected individuals and to maintain population abundance. After sampling and replacement, (1) three liters of *D. magna* growth medium at 5% of the recommended SeO_2_ (ADaM (Klüttgen et al., 1994)) were removed from each experimental population and replaced with three liters of new ADaM to refresh the growth medium, resulting in a *per capita* spore mortality rate of *O. colligata* of γ = 0.0286 days^-1^ (assuming that the spores were well mixed) (Kirk et al., 2020b), and (2) each population was fed 350 million batch cultured algae (*Monoraphidium minutum*).

Populations 1–4 were designated as controls and maintained at a constant temperature of 10°C, which is too cold for the parasite to establish and spread (Fig. 1–2). Populations 5–8 were assigned to a warming treatment, wherein the temperature of the system was set to 10°C at the beginning of the experiment, and increased by 0.5°C every fifteen days, to a maximum of 13.5°C after 105 days. All experiments were terminated after 120 days (Fig. 1–2). Kirk et al.’s (2020) trait-based mechanistic model predicts that R_0_ will exceed the critical threshold of R_0_ = 1 at 12°C, pushing the system through a critical transition from a sub-epidemic state to an epidemic state. As such, only warming populations were subjected to epidemic emergence. This prediction is supported by the experimental data (Kirk et al., 2020b).

**Figure 1:**
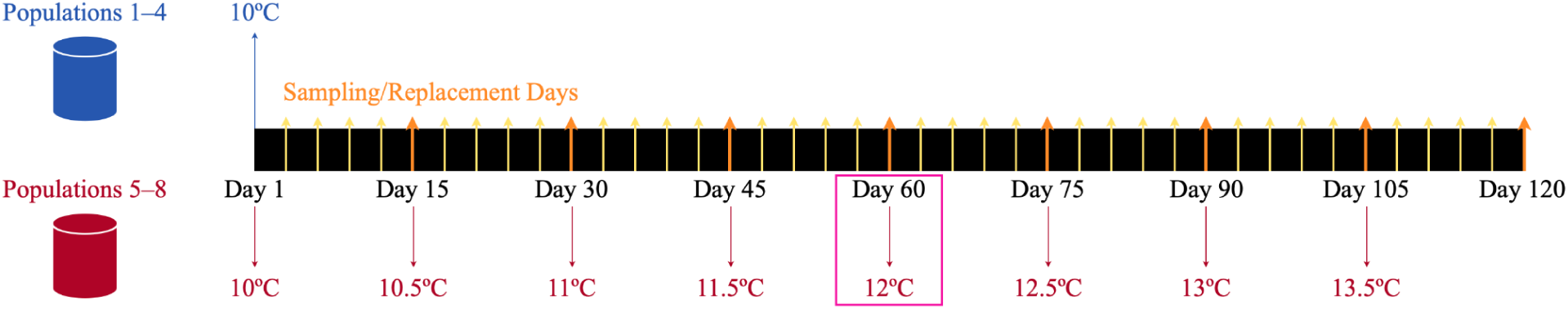
Schematic detailing experimental design. Populations 1–4 were held at 10°C for the duration of the experiment (shown in blue). Populations 5–8 were subjected to a gradual warming treatment (shown in red). Every third day of the experiment, twelve individuals were sampled from each population, and replaced by three randomly selected individuals from infected stock populations, and nine individuals from uninfected stock populations. Sampled individuals were assessed for infection. The pink box represents the day on which the temperature of the system was increased to 12°C, allowing R_0_ to surpass the critical threshold, pushing warming populations from a sub-epidemic state to an epidemic state.

**Figure 2:**
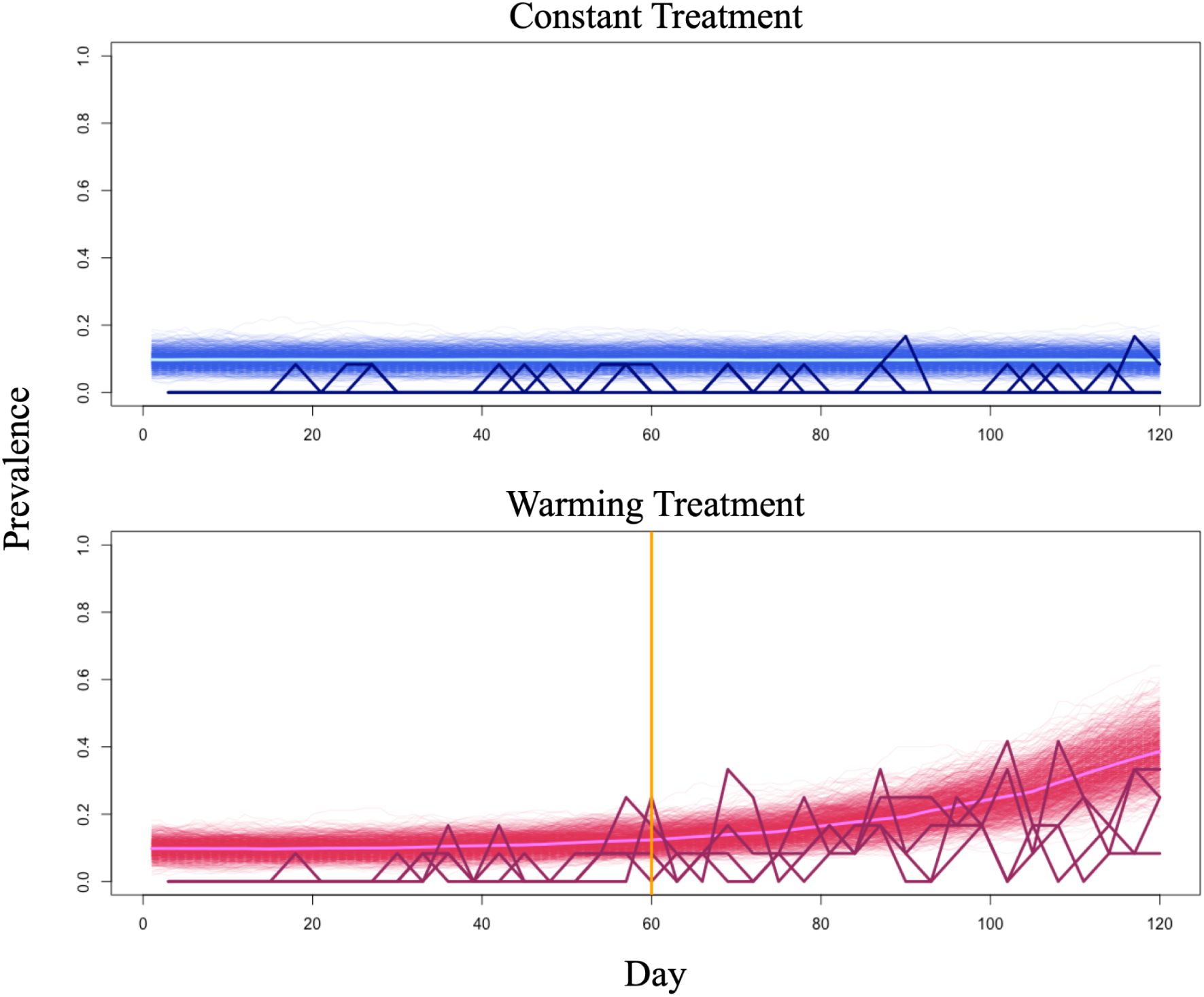
Empirical and simulated time series. In both plots, simulated time series are shown in a lighter shade, while experimental time series are shown in a darker shade. Mean simulated prevalence is shown in the lightest shade. The orange vertical line denotes the day on which the temperature of the system was increased to 12°C, allowing R_0_ to surpass the critical threshold, pushing warming populations from a sub-epidemic state to an epidemic state.

### Model Description

The mechanistic model (Eqns. (1.1)–(1.4)) contains thirteen parameters, six of which are temperature-dependent (*T*), and describes how susceptible hosts (*S*) transition into infecteds (*I*), and subsequently, dead infecteds (*D*) (Table 1). Infecteds (*I*) and dead infecteds (*D*) shed parasite spores into the environment (*E*) (Table 1). All parameters and temperature-dependencies were set as determined by Kirk et al. (2018) and Kirk et al. (2019) (Kirk et al., 2018, 2019, 2020b). For a detailed description of model parameterization and model assumptions, please refer to Kirk et al.’s (2020) Supplementary Material (Kirk et al., 2020a). As per the mechanistic model, susceptible hosts (*S*) are added to each population by the experimenter and as described above (ϕ_*s*_) or through density-dependent recruitment (wherein ψ represents maximum recruitment rate and *K* represents carrying capacity), and lost via infection (at a rate of χ(*T*)σ(*T*)*SE*), natural mortality (µ), or harvesting (ℎ). Infected adults (*I*) are added via immigration (ϕ_*I*_) and transmission (χ(*T*)σ(*T*)*SE*), and lost via natural and parasite-induced mortality (µ(*T*); α(*T*)), and harvesting (ℎ). Dead infecteds (*D*) are added via natural and parasite-induced mortality (µ(*T*); α(*T*)), and are lost via degradation (θ). Spores are released into the environment (*E*) via shedding from living and deceased infecteds (λ(*T*); ω(*T*)) and lost via experimentally induced mortality (γ).

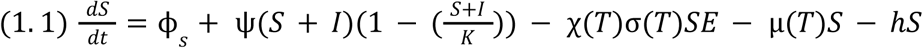

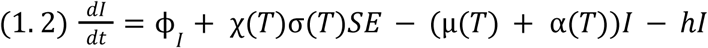

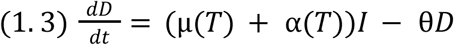

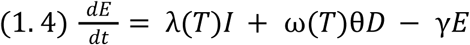

**Table 1:** Parameter definitions and values for the mechanistic *D. magna*–*O. colligata* model (Eqns. (1.1)–(1.4)), replicated from Kirk et al. (2020). Note that the last six parameters are temperature-dependent and are written as functions of temperature, *T*.

| Parameter | Description | Source | Function | Value |
| --- | --- | --- | --- | --- |
| $\phi_s$ | Total rate of input of susceptibles | Kirk et al. (2020) | Constant | 3.535 day<sup>-1</sup> |
| $\phi_I$ | Total rate of input of infecteds | Kirk et al. (2020) | Constant | 0.465 day<sup>-1</sup> |
| $K$ | Adult density-dependent recruitment constraint | Kirk et al. (2020) | Constant | 170 |
| $\psi$ | Maximum recruitment | Kirk et al. (2020) | Constant | 1.33 day<sup>-1</sup> |
| $h$ | Harvesting | Kirk et al. (2020) | Constant | 0.0235 day<sup>-1</sup> |
| $\gamma$ | Environmental spore mortality | Kirk et al. (2020) | Constant | 0.0286 day<sup>-1</sup> |
| $\theta$ | Corpse degradation | Kirk et al. (2020) | Constant | 0.1 day<sup>-1</sup> |
| $\mu(T)$ | Natural mortality rate | Kirk et al. (2018) | Sharpe-Schoolfield<sup>a</sup> | day<sup>-1</sup> |
| $\chi(T)$ | Contact rate | Kirk et al. (2019) | Sharpe-Schoolfield<sup>a</sup> | day<sup>-1</sup> |
| $\sigma(T)$ | Probability of infection | Kirk et al. (2019) | Sharpe-Schoolfield<sup>a</sup> | – |
| $\lambda(T)$ | Parasite shedding rate | Kirk et al. (2018) | Sharpe-Schoolfield<sup>a</sup> | day<sup>-1</sup> |
| $\alpha(T)$ | Parasite-induced mortality rate | Kirk et al. (2018) | Sharpe-Schoolfield<sup>a</sup> | day<sup>-1</sup> |
| $\omega(T)$ | Parasite intensity at host death | Kirk et al. (2018) | Sharpe-Schoolfield<sup>a</sup> | – |

<sup>a</sup>See Figure 1 in Kirk et al. (2020) for functional forms.

### Simulation Experiment

To generate the simulated data, we conducted stochastic simulations of the mechanistic model that matched the duration and environmental conditions of the experimental system. Under constant conditions, the temperature of the system was set to 10°C for the duration of the simulation. Under warming conditions, the temperature of the system was set to 10°C, and increased by 0.5°C every fifteen days. A maximum temperature of 13.5°C was reached after 105 days, and the experiment was terminated after 120 days (Fig. 1). We generated one thousand time series under constant conditions, and one thousand time series under warming conditions (Fig. 2). Demographic stochasticity was introduced via the Gillespie algorithm using the GillespieSSA package in R version 4.2.2 (Pineda-Krch, 2008). Neither the simulations nor the experiment included environmental stochasticity.

### Analysis

#### Statistical Metrics

We considered ten EWS of the transcritical bifurcation at R_0_ = 1 (Table 2). In addition to commonly studied statistics (e.g. variance and autocorrelation, which are expected to increase as a system approaches a local bifurcation point), we considered mean prevalence, which we expected to increase as R_0_ increased towards one (Brett et al., 2018; Harris et al., 2020). We also considered: the coefficient of variation, which relates mean and standard deviation; the index of dispersion, which relates mean and variance; first difference variance; autocovariance, which is the covariance of the process with itself and is related to variance and autocorrelation; and decay time, a log-transform of autocorrelation (Table 2) (Brett et al., 2018; Harris et al., 2020). We also considered two higher-order moments, skewness and kurtosis. Skewness is a measure of the symmetry of the frequency-distribution curve, and kurtosis is a measure of the sharpness of the peak of the frequency-distribution curve (Table 2).

**Table 2:**
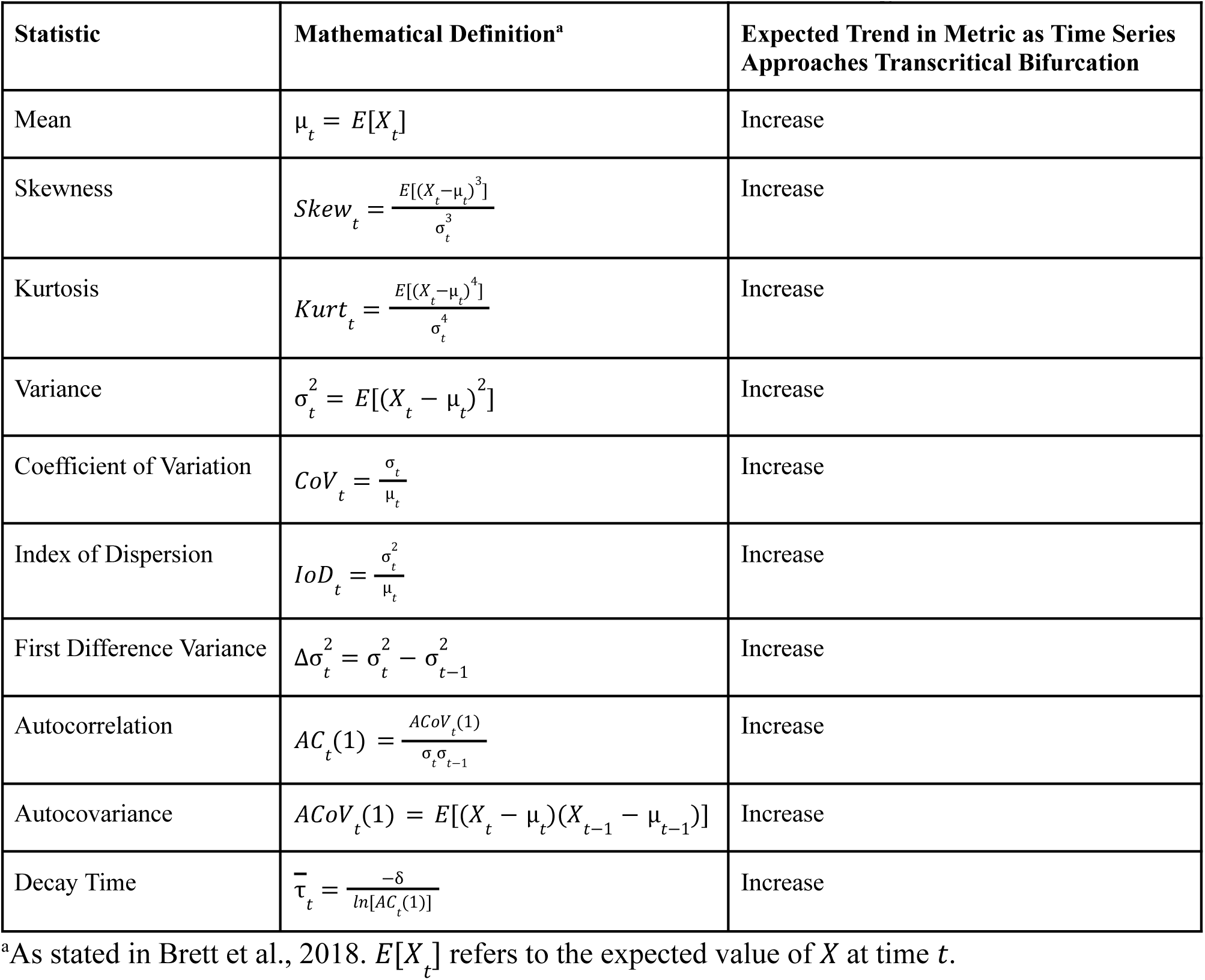
Statistical metrics used to evaluate time series for critical slowing down.

#### Pre-Processing Time Series Data

While it is common practice to pre-process time series data to remove seasonal and longer-term trends (Gama Dessavre et al., 2019; Southall et al., 2021), neither our experimental nor our simulated data were influenced by seasonal forcing, and were collected/generated over relatively short time periods. As such, “detrending” these data may introduce artificial patterns rather than remove them (Delecroix et al., 2022; Miller et al., 2017). In the interest of robustness, we ran analyses on raw and pre-processed data (with pre-processing done using a Gaussian kernel (Dakos et al., 2008, 2012; Harris et al., 2020; O’Regan & Drake, 2013; Pananos et al., 2017; Southall et al., 2021)). We found that pre-processing inflated the strength and reversed the directionality of observed trends, suggesting that pre-processing introduced artificial patterns (Miller et al., 2017). (See supplementary material (Fig. S1–S4) for a more detailed description of these methods and their results.) As such, we will only present and discuss the results of analyses on raw data.

#### Sliding Windows and Kendall’s Tau

We used the ‘evobiR’ package (Heath Blackmon and Richard H. Adams, 2013) in R version 4.2.2 to calculate mean, standard deviation, skewness, and kurtosis. We used the ‘stats’ package in R version 4.2.2 to calculate autocorrelation and autocovariance, and calculated the coefficient of variation, index of dispersion, and first difference variance according to their mathematical definitions, as stated in Table 2 (Brett et al., 2018). All statistical metrics were calculated within sliding windows. We analyzed the simulated data within five-, fifteen-, and thirty-day sliding windows. Given that experimental sampling occurred every three days over a period of 120 days, resulting in forty data points, we were unable to apply a five-day sliding window, and did not have enough data to apply a thirty-day sliding window. As such, we analyzed the empirical data within fifteen-day sliding windows. To evaluate trends in statistics during the approach to criticality, we calculated Kendall’s rank correlation coefficient (or, the trend coefficient). Given that the temperature of the system increased to 12°C on day sixty, we considered three pre-critical intervals: days one to sixty (sixty days), days twenty to sixty (forty days), and days thirty to sixty (thirty days). Analyses of simulated time series were not substantially affected by pre-critical interval choice, while data quality precluded the reliable analysis of empirical time series within the thirty- and forty-day pre-critical intervals. As such, we present analyses of empirical time series performed within the sixty-day pre-critical interval in the main text. (See supplementary material for the full set of results.) Kendall’s τ ({τ ∈ *R* | − 1 ≤ τ ≤ 1}) is a measure of the ordinal association between two objects (Chen et al., 2022). Here, Kendall’s τ is a measure of the association between the rank order of the observed value and its position in time. As such, negative values indicate a decreasing trend prior to local bifurcation, while positive values indicate an increasing trend prior to local bifurcation.

#### Assessing Significance of Empirical Detections of Early Warning Signals

After analyzing the empirical data, we assessed the significance of our detections in four ways. In each case, we used the first half (days one to sixty) of each time series (in warming populations, the pre-bifurcation time series) to generate null distributions of the trend coefficient, against which we compared the true value of the trend coefficient.

First, we used the first half of each time series to parameterize an autoregressive model (AR-1) (Eqn. (2.1)) and simulate one thousand surrogate time series per experimental population (Dakos et al., 2008; Theiler et al., 1992). Within the autoregressive model, β1 is the lag-1 autocorrelation of the residual time series, β_0_ is equal to µ(1 − β_1_) where µ is the mean of the time series, and σ is determined via *v*, the variance of the time series, such that 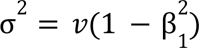 ε_*t*_ is uncorrelated Gaussian noise. We then analyzed the surrogate time series as described above and summarized the resulting trend coefficients as null distributions. We calculated the probability of observing the “true” trend coefficients by chance as the proportion of surrogate trend coefficients equal to or more extreme than the “true” value (Dakos et al., 2008).

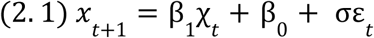

Second, we used the first half of each time series to parameterize an autoregressive model (AR-1), which we combined with a Poisson process to simulate one thousand discrete-valued surrogate time series per population. We used ‘rstan’ (Guo et al., 2015) to estimate β1, µ, and σ such that µ_*t*_ ∼*Normal*(β_1_ · µ_*t*−1_, σ), and the number of infected individuals was Poisson distributed such that 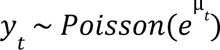 (Fig. S5). To fit the model to each experimental time series, we ran four Markov chains for four thousand iterations (Fig. S5). In all cases, resulting 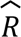 statistics indicated model convergence (Fig. S5). We then analyzed the surrogate time series, summarized the resulting trend coefficients as null distributions, and calculated the probability of observing the “true” trend coefficients by chance as described above.

Third, we used an alternative method of calculating the trend coefficient described by Hamed and Rao (1998) to generate a null distribution. We generated each null distribution by drawing one thousand samples from a normal distribution with a mean of zero and variance given by Eqn 2.2, where *n* is the number of observations in the first half of each time series, and ρ_*s*_ (*i*) is the autocorrelation of the ranks of the observed statistic (e.g. mean, variance, index of dispersion, etc.) (Chen et al., 2022; Hamed & Ramachandra Rao, 1998). While the null hypothesis of the traditional Mann-Kendall trend test is that data are independent, randomly ordered, and without serial correlation, the null hypothesis of Hamed and Rao’s (1998) modified Mann-Kendall trend test is that data do not follow a trend, but are autocorrelated (Chen et al., 2022). Given that most empirical data are autocorrelated, the modified Mann-Kendall trend test improves on its predecessor by decreasing the incidence of false positives. We calculated the probability of observing the “true” trend coefficients by chance as described above.

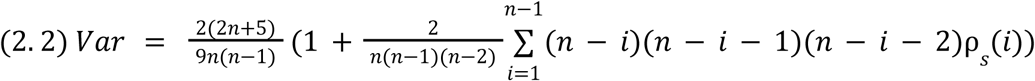

Fourth, we applied a bootstrapping method to the first half of each time series to generate one thousand bootstrapped time series per experimental population (Dakos et al., 2008; Harris et al., 2020). We analyzed each group of bootstrapped time series as described above, and summarized the resulting trend coefficients as null distributions. We calculated the probability of observing the “true” trend coefficients by chance as the proportion of bootstrapped coefficients equal to or more extreme than the “true” value (Dakos et al., 2008; Harris et al., 2020). Within a population, a trend in an indicator was deemed strong if the “true” trend coefficient was equal to or more extreme than 95% of null trend coefficients, across all four testing methods.

Finally, we compared control (constant temperature/non-epidemic) and warming (warming treatment/epidemic emergence) coefficients across simulations and experimental populations by calculating the area under the curve (AUC) statistic. The AUC statistic measures the overlap between null and warming trend coefficient distributions and can be interpreted as the probability that a given warming coefficient will be higher than a given null coefficient (Brett et al., 2018). As such, values less than 0.5 suggest that a decrease in the statistical metric indicates emergence, while values greater than 0.5 suggest that an increase in the statistical metric indicates emergence, with more extreme values indicating stronger trends.

#### Sampling from Simulated Data

After analyzing the simulated data, we conducted a secondary analysis to replicate and account for the effects of experimental sampling. To do so, we (1) removed non-sampling days from each simulated time series, leaving us with prevalence counts for every third day, (2) implemented sampling via a binomial process, such that on every remaining day, twelve samples were taken from each population, with probability of infection given by the number of infecteds divided by total population size (N = 170), and (3) re-ran analyses as described above.

## Results

### Warming Experiment

The empirical data revealed that during the approach to criticality (days one to sixty), both control and warming populations exhibited increases in mean prevalence and variance (Table 3, Fig. 3–4). One control population (four) and one warming population (seven) exhibited increases in mean prevalence (0.000 ≤ *p* ≤ 0.055), while one control population (four) and two warming populations (seven and eight) exhibited increases in variance (0.002 ≤ *p* ≤ 0.05) (Table 4, Fig. 5). With reference to the analysis of mean prevalence, population four exhibited a trend coefficient more extreme than 95% of null trend coefficients across three of four tests, and more extreme than 94.5% of null trend coefficients produced by the remaining test (*P*-value = 0.055). As such, although increasing mean prevalence and variance appeared to precede epidemic emergence within warming populations, the detection of these trends was inconsistent across warming populations and constituted a false positive in a subset of control populations.

**Figure 3:**
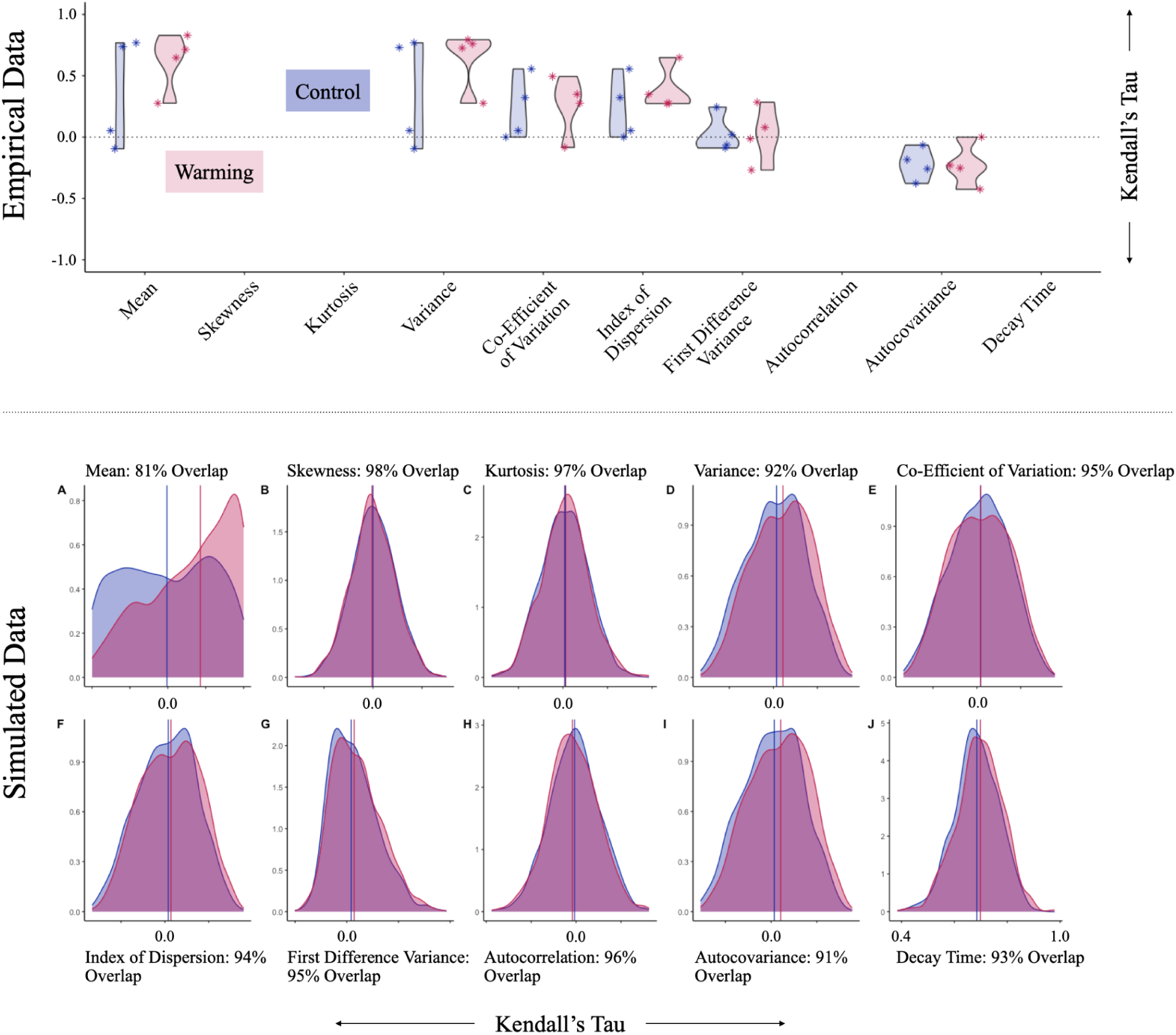
Density and violin plots showing trend coefficients resulting from analysis of empirical and simulated data within a fifteen-day sliding window, between days one and sixty. In all plots, blue represents “Control” results and pink represents “Warming” results. Density plots show the results of one thousand simulations under constant conditions, and one thousand simulations under warming conditions (window size = fifteen days). Density is shown on the vertical axis and trend coefficients are shown on the horizontal axis. Vertical lines represent medians. Violin plots show the trend coefficients resulting from analysis of four populations (1–4) maintained under constant conditions, and four populations (5–8) subjected to a warming treatment. Trend coefficients are shown on the vertical axis and EWS are shown on the horizontal axis.

**Figure 4:**
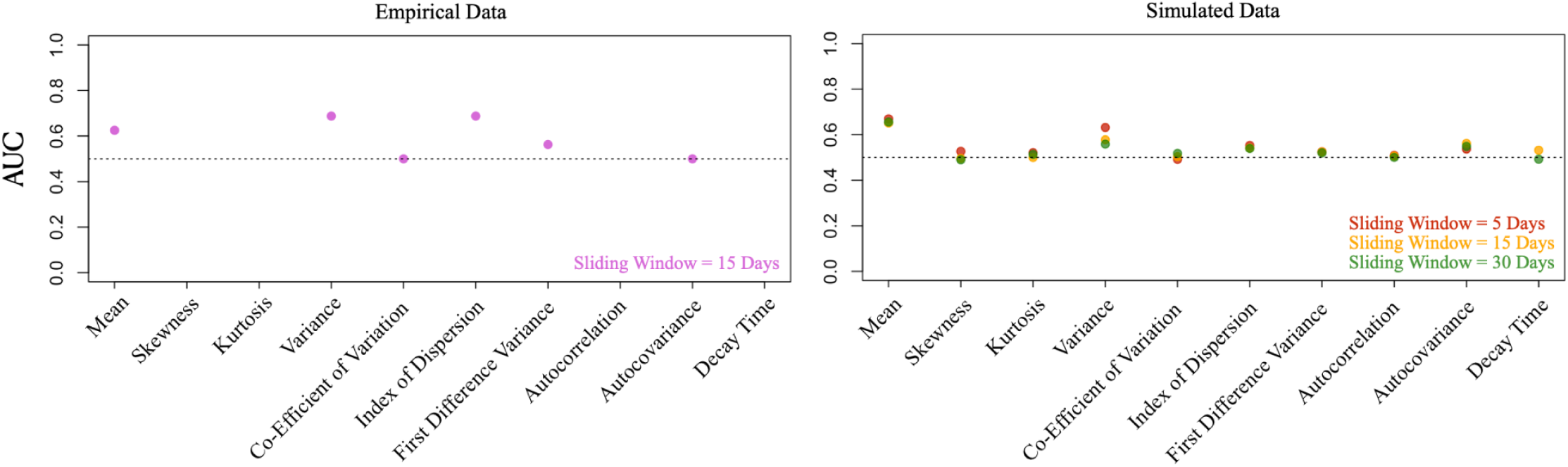
AUC statistics comparing statistical trends in control and test populations. To evaluate statistical trends, we calculated Kendall’s rank correlation coefficient during the pre-critical interval (here, days one to sixty), and compared control (constant temperature/non-epidemic) and warming (warming treatment/epidemic emergence) coefficients across simulations and experimental populations by calculating the area under the curve (AUC) statistic. Values less than 0.5 suggest that a decrease in the statistical metric indicates emergence, while values greater than 0.5 suggest that an increase in the statistical metric indicates emergence, with more extreme values indicating stronger trends. AUC statistics are shown on the vertical axis. EWS are shown on the horizontal axis. Sliding window sizes used to calculate statistical metrics are denoted in the bottom-right of each plot.

**Figure 5:**
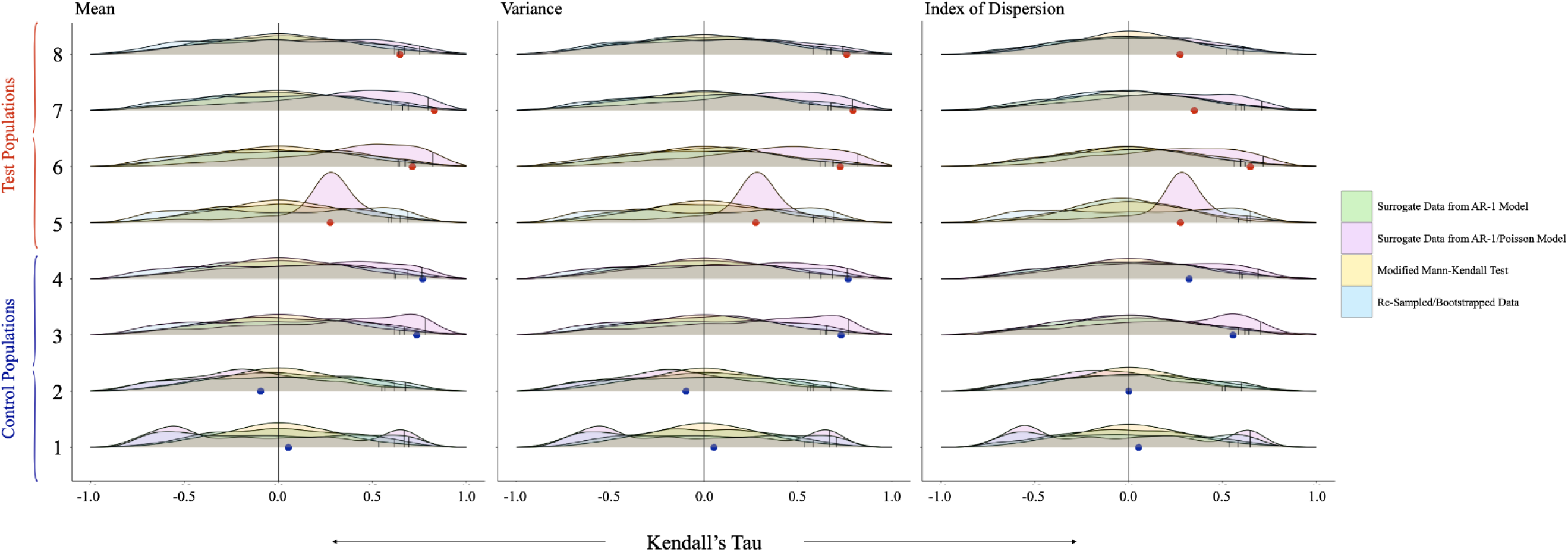
Null distributions to assess detectability of EWS of epidemic emergence in experimental populations. “Control” (Populations 1–4) and “Warming” (Populations 5–8) results are differentiated on the vertical axis. Kendall’s Tau values are shown on the horizontal axis. Green density plots show one thousand Kendall’s Tau values resulting from the simulation of an AR-1 model. Pink density plots show one thousand Kendall’s Tau values resulting from the simulation of an AR-1 model with a Poisson process. Yellow density plots show one thousand Kendall’s Tau values resulting from the application of a modified Mann-Kendall test. Blue density plots show one thousand Kendall’s Tau values resulting from re-sampled time series. Small black vertical ticks represent the 95th quantiles (four ticks for each population, with one tick for each of four tests), while blue (“Control”) and red (“Warming”) dots show true values resulting from the empirical data. Labels above each set of density plots denote the EWS being evaluated.

**Table 3:** Trend coefficients and AUC statistics^a^ as calculated from empirical time series during the approach to criticality (Days 1 to 60). Each statistical metric was calculated within sliding windows, throughout the pre-critical interval. We considered five-, fifteen-, and thirty-day sliding windows. Given that the temperature of the system increased to 12°C on day sixty, we also considered three pre-critical intervals: Days 1 to 60, Days 20 to 60, and Days 30 to 60. To evaluate trends in these metrics, we calculated Kendall’s rank correlation coefficient during the pre-critical interval. Negative values indicate a decreasing trend prior to local bifurcation, while positive values indicate an increasing trend prior to local bifurcation. We compared control (constant temperature/non-epidemic) and warming (warming treatment/epidemic emergence) coefficients across simulations and experimental populations by calculating the area under the curve (AUC) statistic. Values less than 0.5 suggest that a decrease in the statistical metric indicates emergence, while values greater than 0.5 suggest that an increase in the statistical metric indicates emergence, with more extreme values indicating stronger trends.

|  | Population 1 | Population 2 | Population 3 | Population 4 | Population 5 | Population 6 | Population 7 | Population 8 | AUC |
| --- | --- | --- | --- | --- | --- | --- | --- | --- | --- |
| Mean | 0.053 | -0.095 | **0.737** | **0.768** | **0.276** | **0.714** | **0.829** | **0.647** | **0.625** |
| Variance | 0.053 | -0.095 | **0.73** | **0.768** | **0.276** | **0.726** | **0.794** | **0.761** | **0.688** |
| Coefficient of Variation | 0.053 | 0 | **0.556** | **0.322** | **0.276** | **0.494** | **0.349** | -0.084 | 0.5 |
| Index of Dispersion | 0.053 | 0 | **0.556** | **0.322** | **0.276** | **0.648** | **0.349** | **0.274** | **0.688** |
| First Difference Variance | -0.063 | -0.089 | **0.244** | 0.019 | 0.078 | **0.284** | -0.014 | **-0.269** | 0.563 |
| Autocovariance | -0.065 | **-0.184** | **-0.378** | **-0.259** | **-0.231** | 0 | **-0.253** | **-0.425** | 0.5 |

<sup>a</sup>Bolded text indicates the detection of a trend.

**Table 4:**
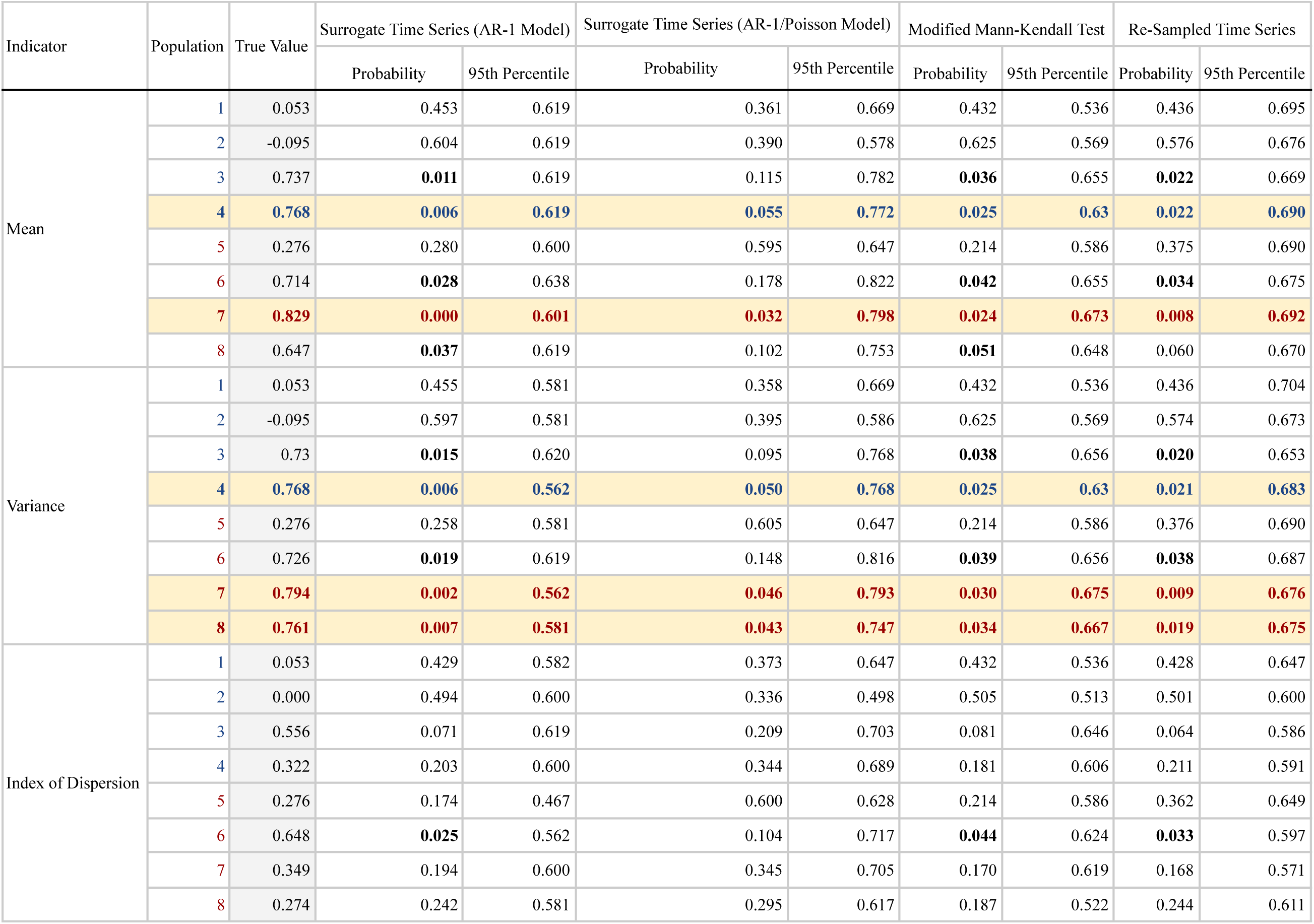
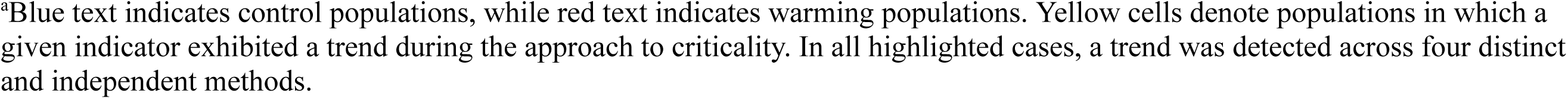
Detectability of EWS of epidemic emergence in experimental populations^a^. We report the true trend coefficient, the proportion of null trend coefficients that are more extreme than the true trend coefficient (“Probability”), and the 95th percentile.

| Indicator | Population | True Value | Surrogate Time Series (AR-1 Model) |  | Surrogate Time Series (AR-1/Poisson Model) |  | Modified Mann-Kendall Test |  | Re-Sampled Time Series |  |
| --- | --- | --- | --- | --- | --- | --- | --- | --- | --- | --- |
|  |  |  | Probability | 95th Percentile | Probability | 95th Percentile | Probability | 95th Percentile | Probability | 95th Percentile |
| Mean | 1 | 0.053 | 0.453 | 0.619 | 0.361 | 0.669 | 0.432 | 0.536 | 0.436 | 0.695 |
|  | 2 | -0.095 | 0.604 | 0.619 | 0.390 | 0.578 | 0.625 | 0.569 | 0.576 | 0.676 |
|  | 3 | 0.737 | **0.011** | 0.619 | 0.115 | 0.782 | **0.036** | 0.655 | **0.022** | 0.669 |
|  | 4 | **0.768** | **0.006** | **0.619** | **0.055** | **0.772** | **0.025** | **0.63** | **0.022** | **0.690** |
|  | 5 | 0.276 | 0.280 | 0.600 | 0.595 | 0.647 | 0.214 | 0.586 | 0.375 | 0.690 |
|  | 6 | 0.714 | **0.028** | 0.638 | 0.178 | 0.822 | **0.042** | 0.655 | **0.034** | 0.675 |
|  | 7 | **0.829** | **0.000** | **0.601** | **0.032** | **0.798** | **0.024** | **0.673** | **0.008** | **0.692** |
|  | 8 | 0.647 | **0.037** | 0.619 | 0.102 | 0.753 | **0.051** | 0.648 | 0.060 | 0.670 |
| Variance | 1 | 0.053 | 0.455 | 0.581 | 0.358 | 0.669 | 0.432 | 0.536 | 0.436 | 0.704 |
|  | 2 | -0.095 | 0.597 | 0.581 | 0.395 | 0.586 | 0.625 | 0.569 | 0.574 | 0.673 |
|  | 3 | 0.73 | **0.015** | 0.620 | 0.095 | 0.768 | **0.038** | 0.656 | **0.020** | 0.653 |
|  | 4 | **0.768** | **0.006** | **0.562** | **0.050** | **0.768** | **0.025** | **0.63** | **0.021** | **0.683** |
|  | 5 | 0.276 | 0.258 | 0.581 | 0.605 | 0.647 | 0.214 | 0.586 | 0.376 | 0.690 |
|  | 6 | 0.726 | **0.019** | 0.619 | 0.148 | 0.816 | **0.039** | 0.656 | **0.038** | 0.687 |
|  | 7 | **0.794** | **0.002** | **0.562** | **0.046** | **0.793** | **0.030** | **0.675** | **0.009** | **0.676** |
|  | 8 | **0.761** | **0.007** | **0.581** | **0.043** | **0.747** | **0.034** | **0.667** | **0.019** | **0.675** |
| Index of Dispersion | 1 | 0.053 | 0.429 | 0.582 | 0.373 | 0.647 | 0.432 | 0.536 | 0.428 | 0.647 |
|  | 2 | 0.000 | 0.494 | 0.600 | 0.336 | 0.498 | 0.505 | 0.513 | 0.501 | 0.600 |
|  | 3 | 0.556 | 0.071 | 0.619 | 0.209 | 0.703 | 0.081 | 0.646 | 0.064 | 0.586 |
|  | 4 | 0.322 | 0.203 | 0.600 | 0.344 | 0.689 | 0.181 | 0.606 | 0.211 | 0.591 |
|  | 5 | 0.276 | 0.174 | 0.467 | 0.600 | 0.628 | 0.214 | 0.586 | 0.362 | 0.649 |
|  | 6 | 0.648 | **0.025** | 0.562 | 0.104 | 0.717 | **0.044** | 0.624 | **0.033** | 0.597 |
|  | 7 | 0.349 | 0.194 | 0.600 | 0.345 | 0.705 | 0.170 | 0.619 | 0.168 | 0.571 |
|  | 8 | 0.274 | 0.242 | 0.581 | 0.295 | 0.617 | 0.187 | 0.522 | 0.244 | 0.611 |

<sup>a</sup>Blue text indicates control populations, while red text indicates warming populations. Yellow cells denote populations in which a given indicator exhibited a trend during the approach to criticality. In all highlighted cases, a trend was detected across four distinct and independent methods.

We did not detect any trends in the coefficient of variation, the index of dispersion, first difference variance, or autocovariance, in any experimental population. Given that empirical time series contained repeated zeros, we were not able to calculate coefficients or AUC statistics for skewness, kurtosis, autocorrelation, and decay time.

### Simulation Experiment

#### Complete Simulated Time Series

Simulated data revealed that as warming populations approached critical transition (R_0_ = 1), mean prevalence and variance increased (Table 5, Fig. 3-4). Across window sizes and pre-critical intervals, analyses of mean prevalence and variance revealed AUC statistics between 0.651 and 0.686, and 0.558 and 0.632, respectively (Table 5, Table S1). We also identified slight increases in the index of dispersion (AUC = 0.518–0.553) and autocovariance (AUC = 0.518–0.562). In all cases, control and warming populations exhibited largely overlapping trend coefficient distributions (mean prevalence: 79% to 81%; variance: 91% to 92%) (Fig. 3, Fig. S7). Different window sizes and pre-critical intervals produced slightly different effect sizes, and identified directionally opposing trends in four indicators: kurtosis, skewness, the coefficient of variation, decay time (Table S1, Fig. S7).

**Table 5:** AUC statistics as calculated from simulated time series. Each statistical metric was calculated within sliding windows, throughout the pre-critical interval. We considered five-, fifteen-, and thirty-day sliding windows. Given that the temperature of the system increased to 12°C on day sixty, we also considered three pre-critical intervals: Days 1 to 60, Days 20 to 60, and Days 30 to 60. To evaluate trends in these metrics, we calculated Kendall’s rank correlation coefficient during the pre-critical interval, and compared control (constant temperature/non-epidemic) and warming (warming treatment/epidemic emergence) coefficients across simulations and experimental populations by calculating the area under the curve (AUC) statistic. Values less than 0.5 suggest that a decrease in the statistical metric indicates emergence, while values greater than 0.5 suggest that an increase in the statistical metric indicates emergence, with more extreme values indicating stronger trends.

| Sliding Window Size | 5 Days |  |  | 15 Days |  |  | 30 Days |
| --- | --- | --- | --- | --- | --- | --- | --- |
| Pre-Critical Interval | Days 1 to 60 | Days 20 to 60 | Days 30 to 60 | Days 1 to 60 | Days 20 to 60 | Days 30 to 60 | Days 1 to 60 |
| Mean | 0.670 | 0.686 | 0.673 | 0.651 | 0.666 | 0.667 | 0.656 |
| Skewness | 0.527 | 0.503 | 0.502 | 0.496 | 0.483 | 0.492 | 0.490 |
| Kurtosis | 0.521 | 0.501 | 0.537 | 0.500 | 0.511 | 0.498 | 0.513 |
| Variance | 0.632 | 0.597 | 0.567 | 0.578 | 0.564 | 0.560 | 0.558 |
| Coefficient of Variation | 0.492 | 0.488 | 0.483 | 0.504 | 0.514 | 0.525 | 0.518 |
| Index of Dispersion | 0.553 | 0.536 | 0.518 | 0.541 | 0.539 | 0.543 | 0.539 |
| First Difference Variance | 0.525 | 0.507 | 0.511 | 0.525 | 0.517 | 0.516 | 0.520 |
| Autocorrelation | 0.509 | 0.516 | 0.503 | 0.506 | 0.513 | 0.523 | 0.500 |
| Autocovariance | 0.537 | 0.531 | 0.518 | 0.562 | 0.549 | 0.549 | 0.549 |
| Decay Time | NA | NA | NA | 0.532 | 0.528 | 0.528 | 0.492 |

#### Subsampled Simulated Time Series

After accounting for the effects of experimental sampling, warming populations continued to exhibit an increase in mean prevalence during the first half of the experiment (days one to sixty) (AUC = 0.622). In contrast to our analysis of the Complete Simulated Time Series, we identified a slight decrease in variance (AUC = 0.463), a slight increase in kurtosis (AUC = 0.560), and slight decreases in the coefficient of variation (AUC = 0.428), the index of dispersion (AUC = 0.440), and autocorrelation (AUC = 0.446) (Table S2, Fig. S8).

## Discussion

We analyzed experimental and simulated time series of disease spread within *Daphnia magna*–*Ordospora colligata* populations for EWS of epidemic emergence. While control populations were maintained at a temperature that prevented epidemic emergence, warming populations were subjected to a warming treatment that forced the system through a critical transition to an epidemic state. We found that EWS of epidemic emergence were nearly as likely to be detected in control populations as they were to be detected in warming populations. Our work constitutes the first experimental study of EWS of epidemic emergence (Dai et al., 2012; Drake & Griffen, 2010; Veraart et al., 2012) and provides theoretical and empirical evidence that EWS may not be a reliable indicator of climate-mediated epidemic emergence.

Our analysis of the empirical data revealed that during the first half of the time series, one control population and one warming population exhibited increases in mean prevalence, and one control population and two warming populations exhibited increases in variance (Tables 3–4, Fig. 3–4). Our analysis of the simulated data revealed that as warming populations approached critical transition, they exhibited increases in mean prevalence and variance (Table 5, Fig. 3–4). We also identified smaller increases in the index of dispersion and autocovariance. However, control and warming populations exhibited largely overlapping trend coefficient distributions (Fig. 3, Fig. S7), suggesting that sub- and pre-epidemic trends may not be easily differentiable. In general, agreement between the empirical and simulated data should increase confidence in our ability to use the trait-based mechanistic model to study temperature-mediated disease emergence in the *Daphnia magna*–*Ordospora colligata* system.

Analyses of the empirical data revealed that EWS of epidemic emergence were nearly as likely to be detected in control populations as they were to be detected in warming populations. We suspect that this outcome is a consequence of the experimental sampling regime and resulting incidence data. First, during the approach to criticality (days one to sixty), samples of control populations one and two identified between zero and one infected individuals, while control populations three and four identified between zero and two infected individuals. Comparatively, samples of warming populations five and six identified between zero and three infected individuals, while samples of warming populations seven and eight identified between zero and five infected individuals. Second, sampling resulted in prevalence counts that oscillated between zero and the maximum detected prevalence (Fig. 2). Despite this, populations four, six, seven, and eight show an increasing trend in detected prevalence (Fig. 2).

Within the simulation experiment, the inclusion of a sampling process revealed that the granularity of prevalence data can drive the magnitude and direction of observed trends (Fig. 4, Fig. S8). While the influence of data quality on the detectability of EWS has been previously discussed (Brett et al., 2018; Clements et al., 2015; Meckawy et al., 2022; O’Brien & Clements, 2021; Proverbio et al., 2022; Southall et al., 2021), our work demonstrates that sampling regimes (i.e. how often sampling occurs, how many samples are collected) are particularly consequential. As such, researchers working with imperfect data (experimental or observational) may benefit from performing power analyses to determine the quality of data required to reliably detect early warning signals. While we did not explicitly consider different types of observation error, additional sources of error would additionally increase the difficulty of detecting EWS.

Given that early warning signals are detected by identifying trends in statistical metrics across sliding windows, small changes and oscillations in prevalence counts are consequential, especially when detected prevalence is low. For example, control populations three and four exhibited a modest jump in detected prevalence from zero to one before day sixty. Despite the fact that both populations exhibited a decrease in detected prevalence after day sixty, this jump results in an increase in mean prevalence, and consequently, the detection of mean prevalence as an early warning signal. Similarly, control populations three and four exhibit their first detection after day twenty. Between first detection and day sixty, both populations experience oscillations in detected prevalence between zero and one, resulting in an increase in variance over time. As such, low and oscillating prevalence counts result in the detection of false positives in control populations, calling into question the reliability of early warning signals detected in warming populations. The impact of data quality on the reliability of EWS is also evident in our analysis of the empirical time series within thirty- and forty-day pre-critical intervals. Given the length and structure of the empirical time series, searching for EWS within smaller pre-critical intervals required us to calculate trend coefficients based on very few data points, exacerbating the impacts of process-independent oscillations, and resulting in unreliable and nonsensical trend coefficients and AUC statistics (Fig. S6).

Data quality is critical to the reliable detection of EWS (Brett et al., 2018; Clements et al., 2015; Meckawy et al., 2022; O’Brien & Clements, 2021; Proverbio et al., 2022; Southall et al., 2021). However, epidemiological data is often imperfect. Specifically, they are often structured by aggregate reporting (Brett et al., 2018) and subject to underreporting, error, and noise (Doyle, 2002; Gibbons et al., 2014; King et al., 2008; O’Dea & Drake, 2019; O’Regan & Burton, 2018). The low and oscillating prevalence counts that limit the utility of our experimental data mirror these issues, and as such, confirm the importance of robust (large and frequent samples in experimental studies to avoid repeated zeros, underestimates of prevalence and process-independent oscillations; carefully collected and pre-processed data in observational studies), long-term time series and speak to the circumstances under which these analyses may result in the detection of false positives (Boettiger & Hastings, 2012; Jäger & Füllsack, 2019). Most importantly, existing works analyse retrospective time series, and are thus unable to use “control” populations to identify false positives. We identified the same EWS of epidemic emergence in control and warming populations, suggesting that false positives may limit the reliability of EWS as predictors of climate-mediated epidemic emergence.

Our simulation experiment, which mimicked a robust sampling regime, revealed largely overlapping trend coefficient distributions (Fig. 3), suggesting that improved sampling may not improve our ability to detect EWS in this system. However, robust and longer-term experimental epidemics in combination with the development of novel methods may still play a role in determining how data quality affects the predictability of epidemic emergence within a range of disease systems (Brett & Rohani, 2020; Bury et al., 2021; O’Regan et al., 2020). In addition, while some systems seem amenable to this approach (Harris et al., 2020; O’Regan et al., 2016), less well-understood systems (Kaur et al., 2020; O’Brien & Clements, 2021; Proverbio et al., 2022) and systems that exhibit transient dynamics (Dablander et al., 2022) can present difficulties. Further, existing works overwhelmingly detect EWS in disease systems characterised by direct transmission (e.g. COVID-19, influenza, pertussis) (Brett et al., 2017; Dablander et al., 2022; Kaur et al., 2020; O’Brien & Clements, 2021; Proverbio et al., 2022; Southall et al., 2021) or vector-borne transmission (e.g. malaria) (Harris et al., 2020; Hussain-Alkhateeb et al., 2021; O’Regan et al., 2016). While the recency of the COVID-19 pandemic and increasing concern over the spread of vector-borne diseases in a warming climate may bias researchers towards studying such systems, our findings suggest that predicting the epidemic emergence of environmentally transmitted diseases may present novel difficulties. Resolving such difficulties will become increasingly important in the face of increasingly frequent extreme climate events which reduce access to clean water, and promote the spread of water-borne diseases like cholera (Charnley et al., 2022).

In summary, we analyzed experimental and simulated time series of disease spread within *Daphnia magna*–*Ordospora colligata* populations for EWS of epidemic emergence. We identified the same EWS of epidemic emergence in control and warming populations, suggesting that false positives confound the interpretation of EWS in pre-epidemic systems. Given that control populations do not exist in natural systems, our findings suggest that EWS may not be a reliable indicator of climate-mediated epidemic emergence. Our findings also identify potential pitfalls in the detection of EWS in disease systems characterised by environmental transmission, suggesting that the detectability of EWS may vary by transmission mode. As our climate changes, our ability to predict climate-mediated epidemic emergence will become more important than ever. Our findings echo the importance of robust and long-term disease surveillance programs, and demonstrate the need for further investigation via experimental epidemics, and into the predictability of differentially transmitted disease outbreaks.

## Acknowledgements

We thank a number of anonymous reviewers, who, through their generous comments, greatly improved the quality of the work.

## Author Contributions

MJ-C, DK, PM, and MK conceived the study. All authors contributed to designing the methodology. DK, LK, and PL designed and conducted the warming experiment. MJ-C and DK analysed the data. MJ-C led the writing of the manuscript. All authors contributed critically to drafts and gave final approval for publication.

