## Supplemental Information for "Early warning signals do not predict a warming-induced experimental epidemic"

**Submitted To: Ecological Applications.**

[1]Madeline Jarvis-Cross (ORCID: 0009-0008-0527-9369)
[2,3]Devin Kirk (ORCID: 0000-0001-9588-1004)
[1]Leila Krichel (ORCID: 0000-0002-0501-3033)
[4]Pepijn Luijckx (ORCID: 0000-0002-8173-6727)
[1,5]Péter Molnár (ORCID: 0000-0001-7260-2674)
[1]Martin Krkosek (ORCID: 0000-0001-7591 7954)

[1]Department of Ecology and Evolutionary Biology, University of Toronto
[2]Department of Biology, Stanford University
[3]Department of Zoology, University of British Columbia
[4]School of Natural Sciences, Trinity College Dublin
[5]Department of Biological Sciences, University of Toronto, Scarborough

#### Table of Contents

Description of data pre-processing 2
Figure S1: Empirical and simulated time series pre-processed with bandwidth of two 3
Figure S2: Empirical and simulated time series pre-processed with bandwidth of three 4
Figure S3: Empirical and simulated time series pre-processed with bandwidth of four 5
Figure S4: AUC statistics as calculated from pre-processed empirical and simulated data 6
Figure S5: Posterior visualizations of autoregressive model with Poisson process 7
Figure S6: Trend coefficients resulting from analysis of experimental time series within different pre-critical intervals 9
Figure S7: Density plots showing trend coefficient distributions within different pre-critical intervals 10
Table S1: Median trend coefficients and AUC statistics as calculated from simulated time series within different sliding windows and pre-critical intervals 11
Table S2: Median trend coefficients and AUC statistics as calculated from simulated time series after accounting for effects of experimental sampling 13
Figure S8: AUC statistics as calculated from simulated data after accounting for the effects of sampling 14

**Description of data pre-processing**

As described in the main text, we used Gaussian detrending to pre-process the simulated and experimental time series. Gaussian detrending, sometimes referred to as Gaussian filtering or Gaussian smoothing, is a moving average technique that involves using a Gaussian kernel (Eqn. 1) to calculate a weighted mean over a selected bandwidth to remove noise and/or non-focal trends (e.g. seasonal trends). In more detail, at every data point $x$, one uses a Gaussian kernel to calculate the kernel function (Eqn. 1), and estimate $x$ as the weighted mean of a specified number of data points (the bandwidth) on either side of $x$ (where the kernel function determines weights. Broadly, kernels define the shape of the function used to take the average of neighboring points, and as such, Gaussian kernels lend a heavier weight to nearby data points. The bandwidth of the kernel defines the number of data points included in the weighted average. As such, narrower kernels risk maintaining unwanted trends, while wider kernels risk removing the signal.

$$(1)\ k(x,\ x_i) = exp(-\frac{||x-x_i||^2}{2b^2})$$

Where $i = 1, ..., M$, and represent the distance between the focal point, $x$, and some other point within the specified bandwidth, $b$[1].

We detrended the simulation and experimental time series using bandwidths of two (Fig. S1), three (Fig. S2), and four (Fig. S3). As shown in Fig. S4, when applied to the experimental data, detrending inflated the strength and/or changed the directionality of observed trends. When applied to the simulated data, detrending had very small effects on the magnitudes of observed statistical trends. Given that pre-processing is expected to decrease the strength of observed trends, that increases in focal metrics should precede a transcritical bifurcation, and that neither experimental nor simulated conditions should have been influenced by seasonality, we suspect that in this case, detrending the empirical and simulated data introduced artificial trends, and thus produced unreliable results.

[1] Xiahai Zhuang and Yipeng Hu, 'Statistical Deformation Model: Theory and Methods', in *Statistical Shape and Deformation Analysis* (Elsevier, 2017), 33–65, https://doi.org/10.1016/B978-0-12-810493-4.00003-1.

**Figure S1: Empirical and simulated time series pre-processed with bandwidth of two**

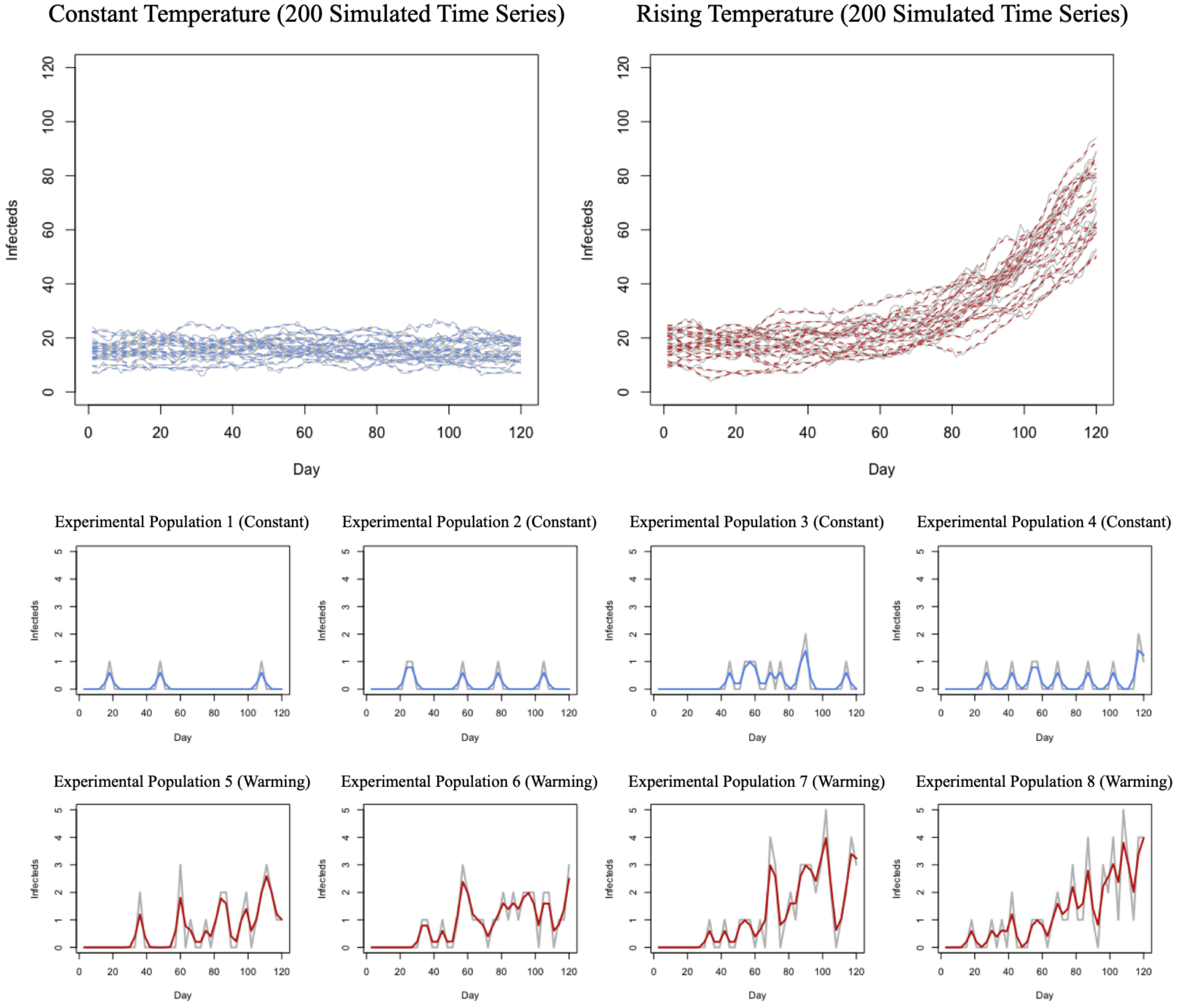


Figure S1: Raw data are shown in grey. Pre-processed time series are shown in blue (control populations) and red (warming populations). These data were pre-processed with a Gaussian kernel, with a bandwidth of two. A subset of 200 simulated time series is shown in each of the top panels; experimental time series are shown in the bottom panels.

**Figure S2: Empirical and simulated time series pre-processed with bandwidth of three**

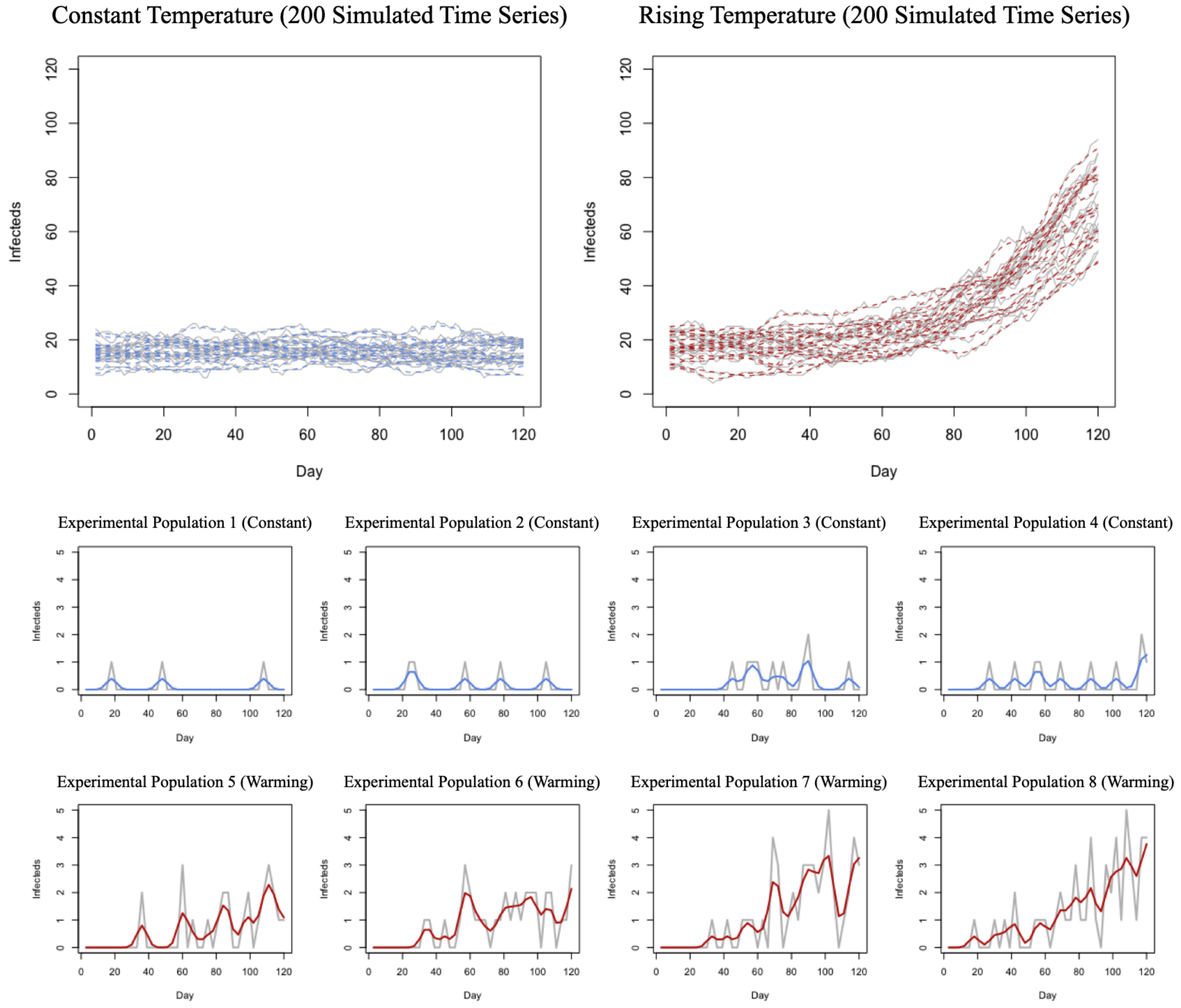


Figure S2: Raw data are shown in grey. Pre-processed time series are shown in blue (control populations) and red (warming populations). These data were pre-processed with a Gaussian kernel, with a bandwidth of three. A subset of 200 simulated time series is shown in each of the top panels; experimental time series are shown in the bottom panels.

#### Figure S3: Empirical and simulated time series pre-processed with bandwidth of four

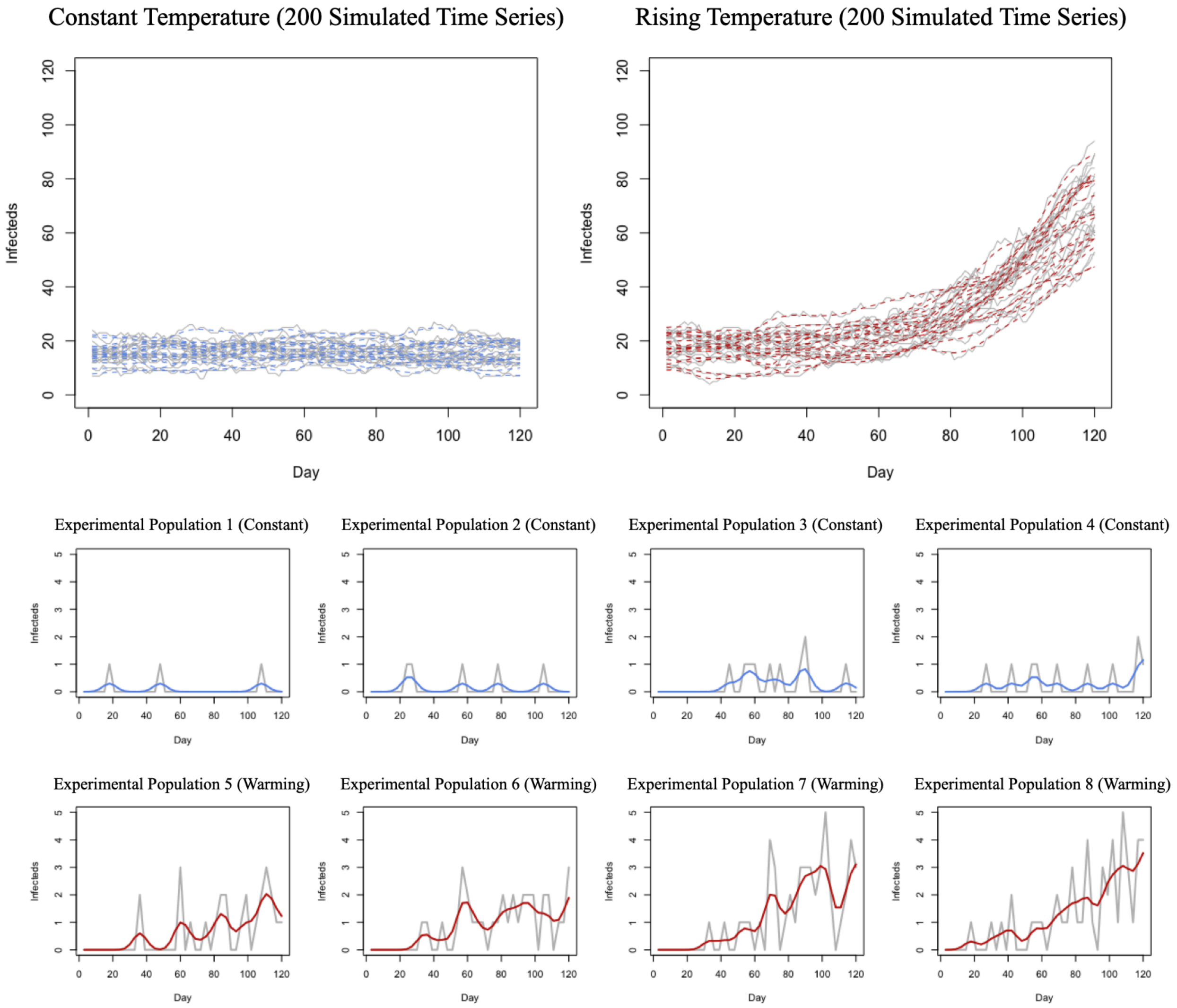


Figure S3: Raw data are shown in grey. Pre-processed time series are shown in blue (control populations) and red (warming populations). These data were pre-processed with a Gaussian kernel, with a bandwidth of four. A subset of 200 simulated time series is shown in each of the top panels; experimental time series are shown in the bottom panels.

#### Figure S4: AUC statistics as calculated from pre-processed empirical and simulated data

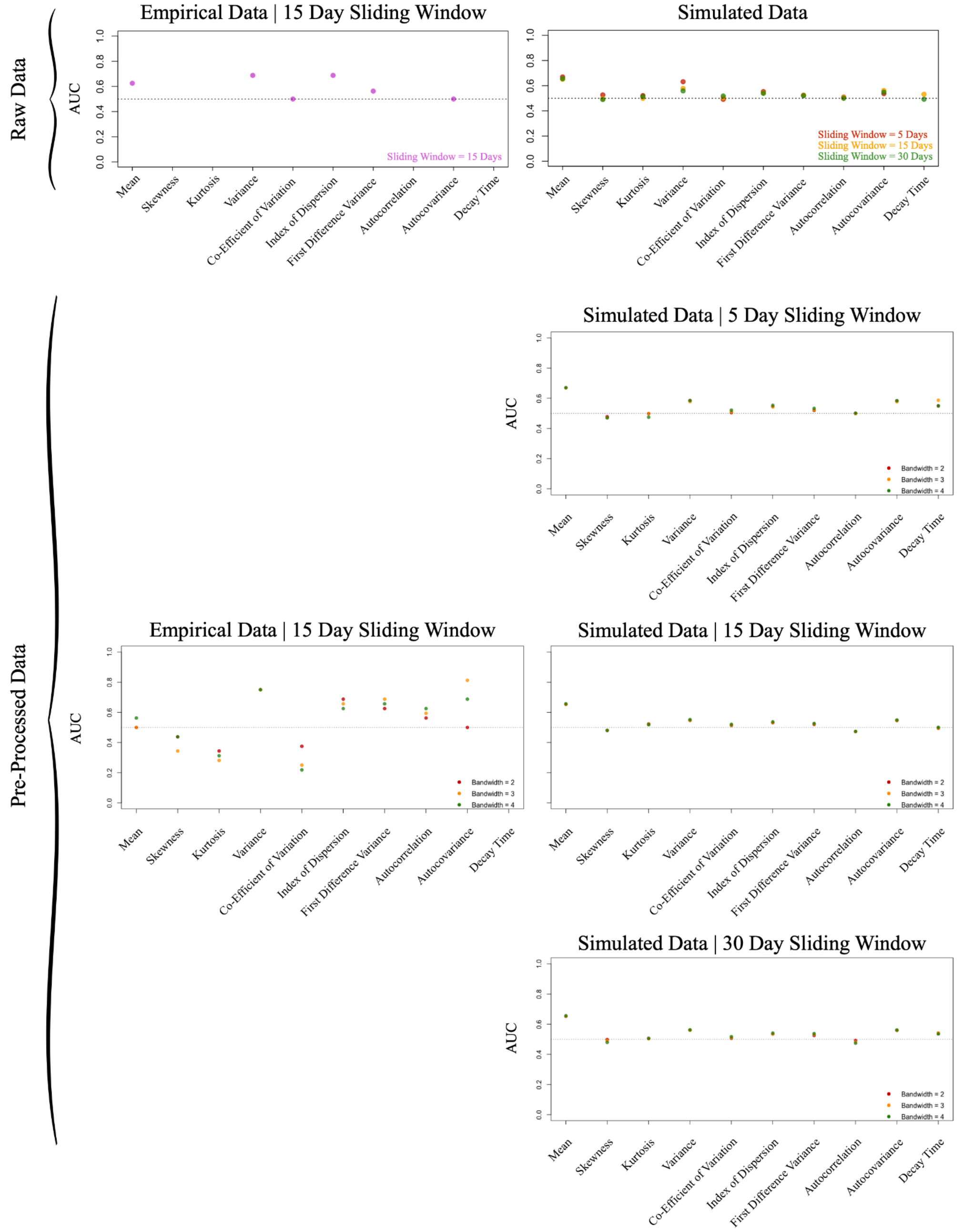


Figure S4: To evaluate statistical trends, we calculated Kendall's rank correlation coefficient during the pre-critical interval (here, Days 1 to 60), and compared control (constant temperature/non-epidemic) and warming (warming treatment/epidemic emergence) coefficients across simulations and experimental populations by calculating the area under the curve (AUC) statistic. Values less than 0.5 suggest that a decrease in the statistical metric indicates emergence, while values greater than 0.5 suggest that an increase in the statistical metric indicates emergence, with more extreme values indicating stronger trends. AUC statistics are shown on the vertical axis. EWS are shown on the horizontal axis. Sliding window sizes used to calculate statistical metrics are denoted in plot titles. Gaussian kernel bandwidths used to pre-process data are denoted in the bottom-right.

### Figure S5: Posterior visualizations of autoregressive model with Poisson process

A

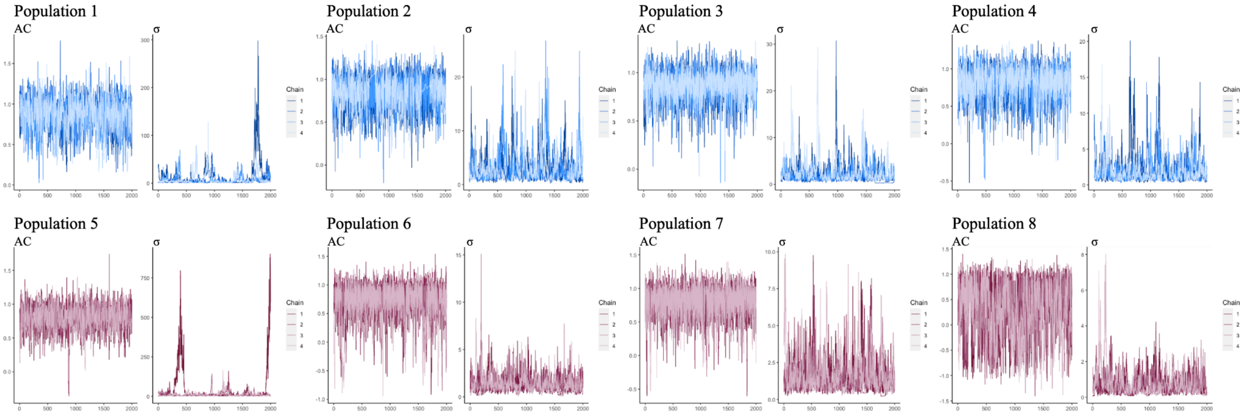


B

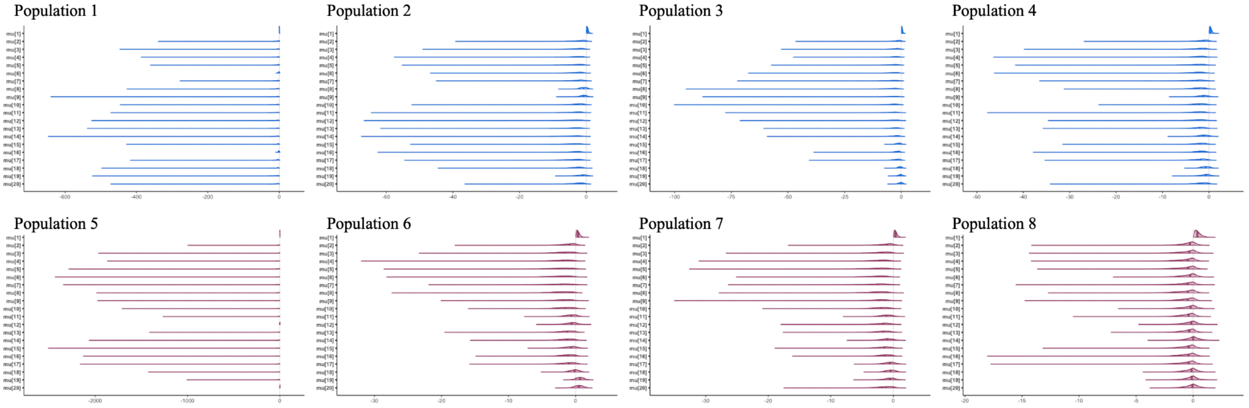


C

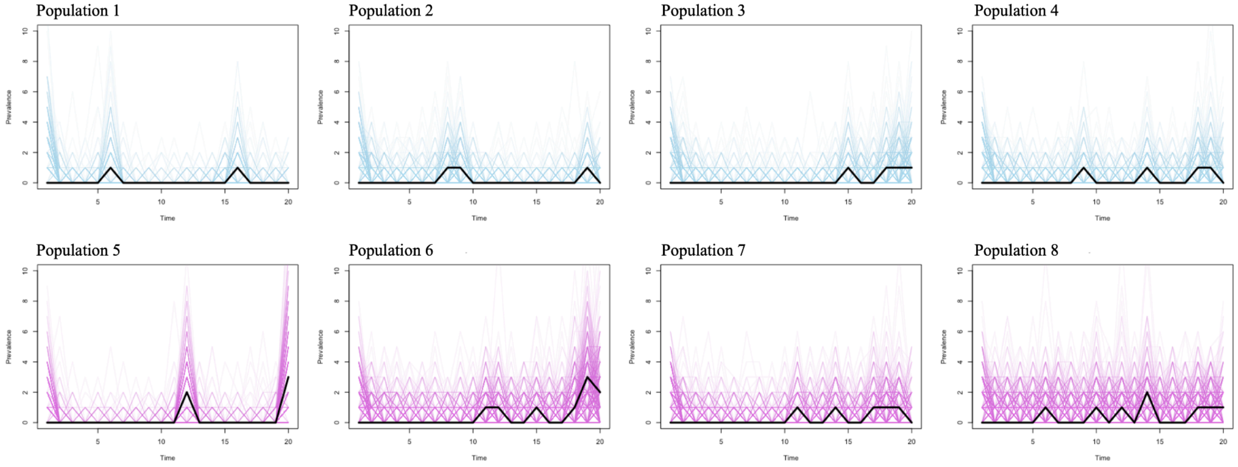


D

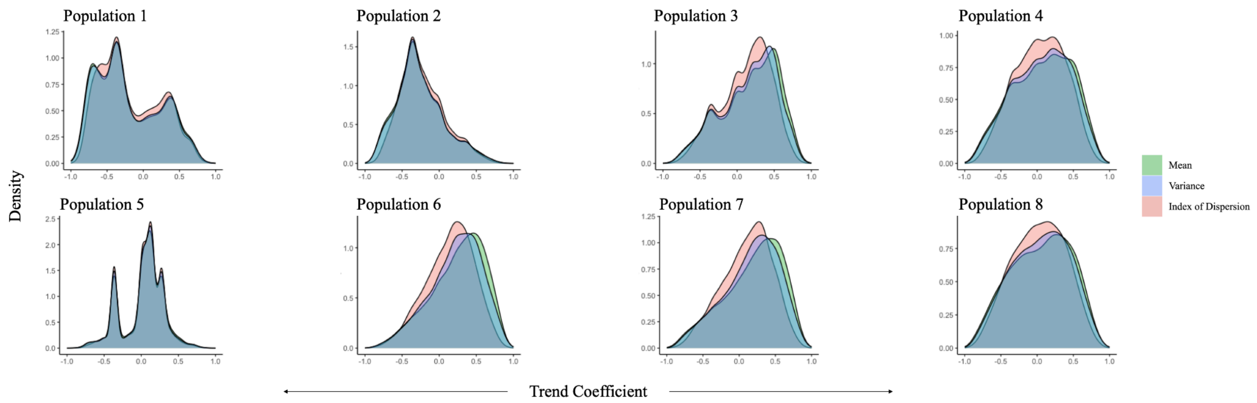


Figure S5: In (A), (B), and (C), control populations are shown in blue, and warming populations are shown in pink. (A) Per the autoregressive model described in the main text (Eqn. (2.1)), trace plots showing sampled values of $\beta_1$ (lag-1 autocorrelation of the residual time series) and σ (variance of the time series) across four chains, over time. (B) Posterior distributions of μ (mean of the time series). (C) Experimental times series overlaid atop model estimated time series that were used to generate a null distribution of trend coefficients. In (D), we show resulting null distributions of trend coefficients.

**Figure S6: Trend coefficients resulting from analysis of experimental time series within different pre-critical intervals**

Days 20 to 60 (15 Day Sliding Window)

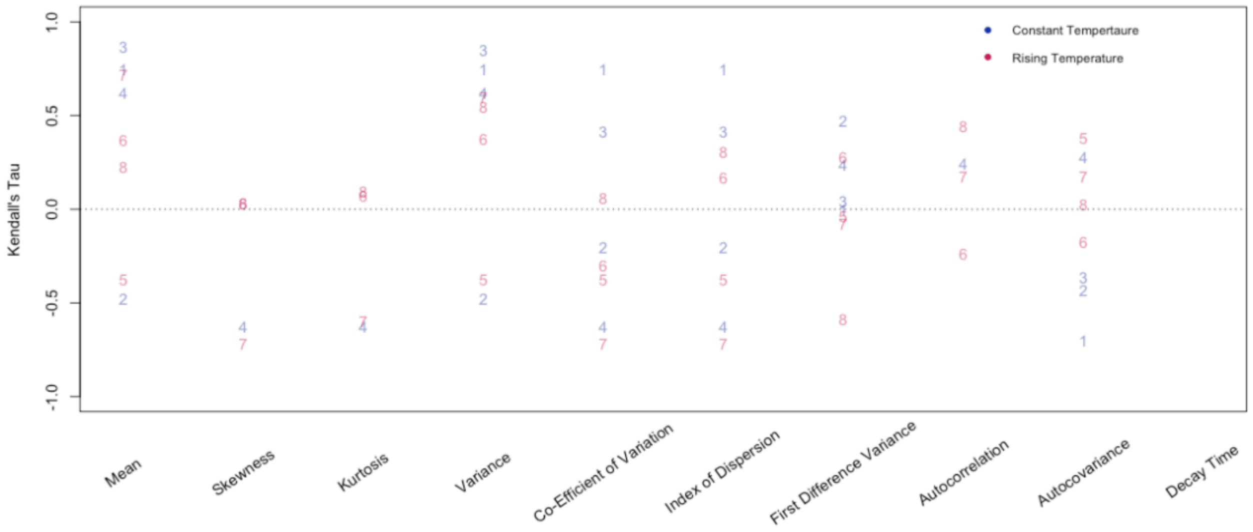


Days 30 to 60 (15 Day Sliding Window)

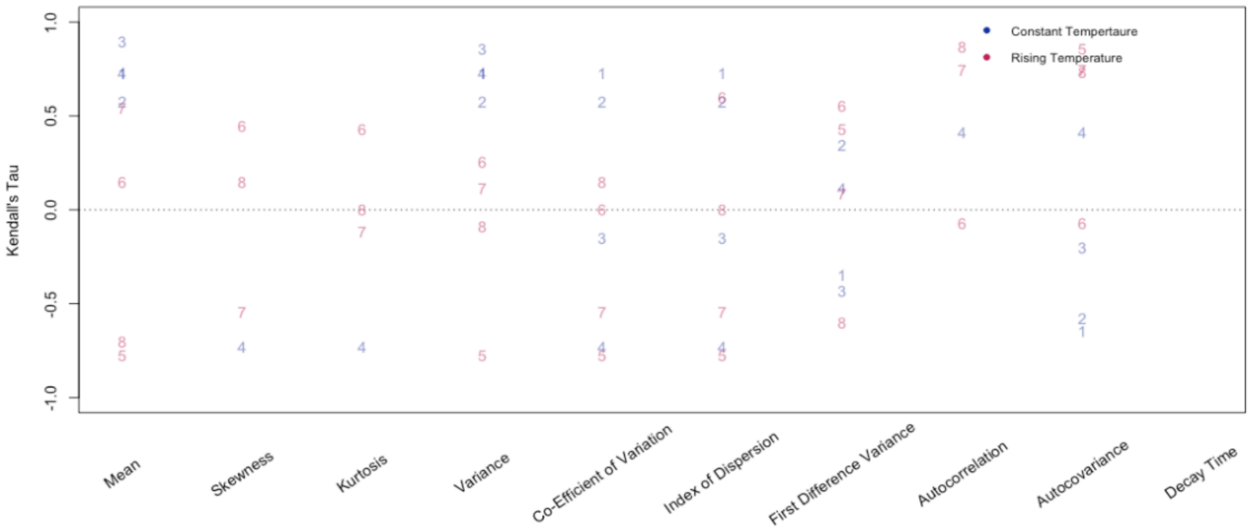


Figure S6: Trend coefficients resulting from analysis of four experimental populations (1–4) maintained under constant conditions (blue), and four experimental populations (5–8) subjected to a warming treatment (pink). Statistical metrics were calculated within fifteen-day sliding windows. To evaluate statistical trends, we calculated Kendall's rank correlation coefficient during the pre-critical interval (forty days (top panel), thirty days (bottom panel), and compared control (constant temperature/non-epidemic) and warming (warming treatment/epidemic emergence) coefficients across simulations and experimental populations by calculating the area under the curve (AUC) statistic. Values less than 0.5 indicate that a decrease in the indicator indicates emergence, while values greater than 0.5 indicate an increasing trend, with more extreme values indicating stronger trends. AUC statistics are shown on the vertical axis. EWS are shown on the horizontal axis.

**Figure S7: Density plots showing trend coefficient distributions within different pre-critical intervals**

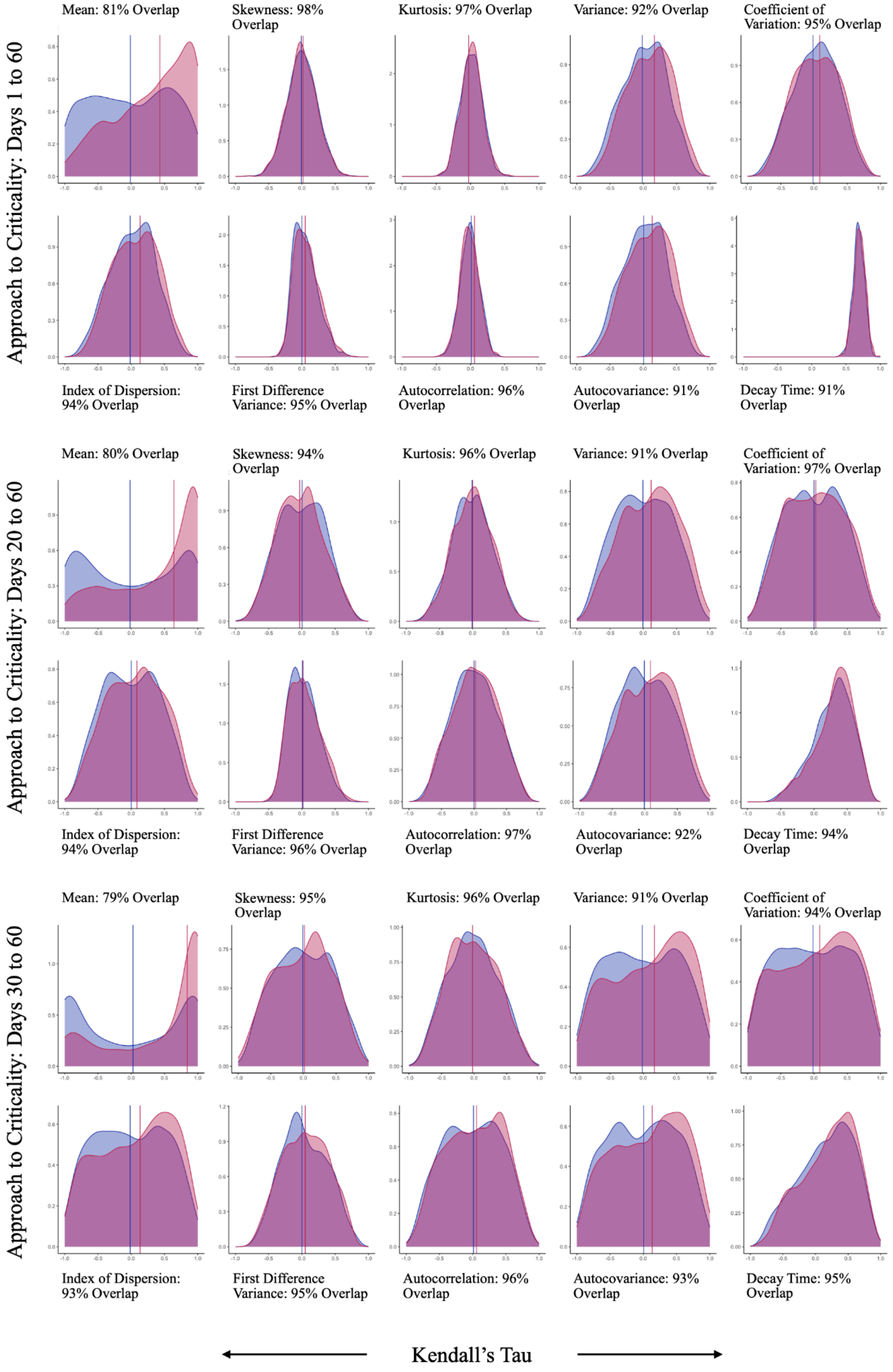


Figure S7: Density plots show the results of one thousand simulations under constant conditions (blue), and one thousand simulations under warming conditions (pink). Density is shown on the vertical axis and trend coefficients are shown on the horizontal axis. Vertical lines represent medians. Statistical metrics were calculated within fifteen-day sliding windows, during sixty-, forty-, and thirty-day pre-critical intervals. The first panel replicates the density plots shown in Fig. 3, for reference.

**Table S1: Median trend coefficients and AUC statistics as calculated from simulated time series[a]. While median tau values are also available in the main text (Table 5), here, we additionally report AUC statistics to directly compare control and warming populations.**

| | Sliding Window: 5 Days | | | | | | Sliding Window: 15 Days | | | | | | Sliding Window: 30 Days | |
|---|---|---|---|---|---|---|---|---|---|---|---|---|---|---|
| | Days 1 to 60 | | Days 20 to 60 | | Days 30 to 60 | | Days 1 to 60 | | Days 20 to 60 | | Days 30 to 60 | | Days 1 to 60 | |
| | Median Tau | AUC | Median Tau | AUC | Median Tau | AUC | Median Tau | AUC | Median Tau | AUC | Median Tau | AUC | Median Tau | AUC |
| Mean | 0.008 | | -0.027 | 0.686 | 0.002 | 0.673 | -0.011 | 0.651 | -0.019 | 0.666 | 0.028 | 0.667 | 0.023 | 0.656 |
| | 0.329 | 0.670 | 0.438 | | 0.474 | | 0.410 | | 0.643 | | 0.843 | | 0.693 | |
| Skewness | 0.004 | | 0.005 | 0.503 | 0.000 | 0.502 | -0.018 | 0.496 | 0.003 | 0.483 | -0.008 | 0.492 | -0.025 | 0.490 |
| | 0.010 | 0.527 | 0.002 | | 0.000 | | -0.015 | | -0.034 | | 0.017 | | -0.046 | |
| Kurtosis | 0.001 | | 0.010 | 0.501 | -0.003 | 0.537 | 0.002 | 0.500 | -0.008 | 0.511 | -0.025 | 0.498 | -0.009 | 0.513 |
| | 0.010 | 0.521 | 0.007 | | 0.013 | | -0.002 | | 0.002 | | -0.025 | | 0.006 | |
| Variance | -0.006 | | -0.004 | 0.597 | -0.010 | 0.567 | 0.013 | 0.578 | -0.006 | 0.564 | -0.017 | 0.560 | 0.016 | 0.558 |
| | 0.084 | 0.632 | 0.075 | | 0.076 | | 0.133 | | 0.118 | | 0.167 | | 0.176 | |
| Coefficient of Variation | -0.004 | | -0.008 | 0.488 | -0.003 | 0.483 | 0.013 | 0.504 | 0.000 | 0.514 | -0.017 | 0.525 | 0.016 | 0.518 |
| | 0.003 | 0.492 | -0.017 | | -0.013 | | 0.015 | | 0.028 | | 0.084 | | 0.092 | |
| Index of Dispersion | 0.003 | | -0.003 | 0.536 | -0.008 | 0.518 | 0.027 | 0.541 | 0.002 | 0.539 | -0.017 | 0.543 | 0.025 | 0.539 |
| | 0.035 | 0.553 | 0.024 | | 0.025 | | 0.087 | | 0.085 | | 0.133 | | 0.145 | |
| First Difference Variance | 0.005 | | 0.005 | 0.507 | 0.002 | 0.511 | 0.007 | 0.525 | 0.007 | 0.517 | 0.000 | 0.516 | 0.058 | 0.520 |
| | 0.010 | 0.525 | 0.005 | | 0.006 | | 0.017 | | 0.019 | | 0.050 | | 0.089 | |
| Autocorrelation | 0.014 | | 0.003 | 0.516 | 0.012 | 0.503 | 0.012 | 0.506 | -0.002 | 0.513 | 0.013 | 0.523 | 0.039 | 0.500 |
| | 0.023 | 0.509 | 0.013 | | -0.006 | | 0.017 | | 0.022 | | 0.059 | | 0.030 | |
| Autocovariance | -0.001 | | 0.005 | 0.531 | 0.000 | 0.518 | 0.018 | 0.562 | -0.006 | 0.549 | 0.004 | 0.549 | 0.023 | 0.549 |
| | 0.019 | 0.537 | 0.019 | | 0.013 | | 0.098 | | 0.088 | | 0.133 | | 0.177 | |
| Decay Time | NA | | NA | NA | NA | NA | 0.610 | 0.532 | 0.323 | 0.528 | 0.217 | 0.528 | 0.784 | 0.492 |
| | NA | NA | NA | | NA | | 0.620 | | 0.354 | | 0.267 | | 0.779 | |

[a]Blue text denotes control populations, while red text denotes warming populations.

Each statistical metric was calculated within five-, fifteen-, and thirty-day sliding windows, during sixty-, forty-, and thirty-day pre-critical intervals. To evaluate trends in these metrics, we calculated a median trend coefficient during the pre-critical interval over one thousand control and one thousand warming time series. Negative values indicate a decreasing trend prior to local bifurcation, while positive values indicate an increasing trend prior to local bifurcation. We compared control (constant temperature/non-epidemic) and warming (warming treatment/epidemic emergence) coefficients across simulations and experimental populations by calculating the area under the curve (AUC) statistic. Values less than 0.5 suggest that a decrease in the statistical metric indicates emergence, while values greater than 0.5 suggest that an increase in the statistical metric indicates emergence, with more extreme values indicating stronger trends.

**Table S2: Median trend coefficients and AUC statistics as calculated from simulated time series after accounting for effects of experimental sampling during the sixty-day pre-critical interval[a].**

| | Sliding Window: 15 Days | |
|---|---|---|
| | Median Tau | AUC |
| Mean | 0.012 | 0.622 |
| | 0.386 | |
| Skewness | 0.021 | 0.503 |
| | -0.031 | |
| Kurtosis | 0.017 | 0.560 |
| | 0.092 | |
| Variance | 0.000 | 0.463 |
| | -0.058 | |
| Coefficient of Variation | 0.000 | 0.428 |
| | -0.131 | |
| Index of Dispersion | 0.000 | 0.440 |
| | -0.096 | |
| First Difference Variance | 0.011 | 0.481 |
| | 0.000 | |
| Autocorrelation | 0.015 | 0.446 |
| | -0.058 | |
| Autocovariance | 0.013 | 0.496 |
| | 0.000 | |
| Decay Time | NA | NA |
| | NA | |

[a]Blue text denotes control populations, while red text denotes warming populations.

To evaluate trends in these metrics, we calculated a median trend coefficient during the pre-critical interval over one thousand control and one thousand warming time series. Negative values indicate a decreasing trend prior to local bifurcation, while positive values indicate an increasing trend prior to local bifurcation. We compared control (constant temperature/non-epidemic) and warming (warming treatment/epidemic emergence) coefficients across simulations and experimental populations by calculating the area under the curve (AUC) statistic. Values less than 0.5 suggest that a decrease in the statistical metric indicates emergence, while values greater than 0.5 suggest that an increase in the statistical metric indicates emergence, with more extreme values indicating stronger trends.

**Figure S8: AUC statistics as calculated from simulated data after accounting for the effects of sampling**

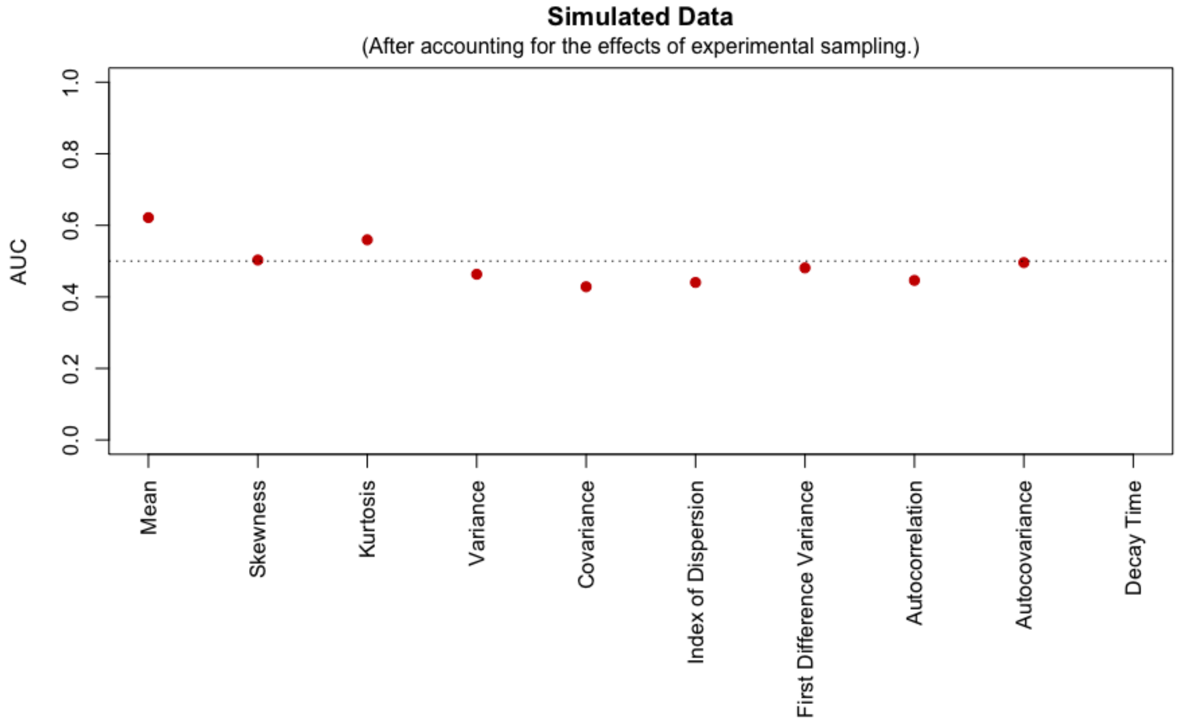


Figure S8: To evaluate statistical trends, we calculated Kendall's rank correlation coefficient during the pre-critical interval, and compared control (constant temperature/non-epidemic) and warming (warming treatment/epidemic emergence) coefficients across simulations and experimental populations by calculating the area under the curve (AUC) statistic. Values less than 0.5 suggest that a decrease in the statistical metric indicates emergence, while values greater than 0.5 suggest that an increase in the statistical metric indicates emergence, with more extreme values indicating stronger trends. AUC statistics are shown on the vertical axis. EWS are shown on the horizontal axis. Analyses were performed within fifteen-day sliding windows, between days one and sixty.
